# Integrated analysis of gene expression and protein-protein interaction with tensor decomposition

**DOI:** 10.1101/2023.02.26.530076

**Authors:** Y-H. Taguchi, Turki Turki

## Abstract

**Motivation:** Integration of gene expression (GE) and protein-protein interaction (PPI) is not straightforward because the former is provided as a matrix, whereas the latter is provided as a network. In many cases, genes processed with GE analysis are refined further based on a PPI network or vice versa. This is hardly regarded as a true integration of GE and PPI. To address this problem, we proposed a tensor decomposition (TD) based method that can integrate GE and PPI prior to any analyses where PPI is also formatted as a matrix to which singular value decomposition (SVD) is applied.

**Results:** Integrated analyses with TD improved the coincidence between vectors attributed to samples and class labels over 27 cancer types retrieved from The Cancer Genome Atlas Program (TCGA) toward five class labels. Enrichment using genes selected with this strategy were also improved with the integration using TD. The PPI network associated with the information on the strength of the PPI can improve the performance than PPI that stores only if the interaction exists in individual pairs. In addition, even restricting genes to the intersection of GE and PPI can improve coincidence and enrichment.

**Availability and implementation:** The R source code used to perform this analyses is in the supplementary file.

## 1. Introduction

Integrated analysis of gene expression (GE) and protein-protein interaction (PPI) has not been studied extensively [1]. This is possibly because of the distinct formats provided for GE and PPI. GE is usually provided in a matrix format, whose columns and rows typically correspond to samples and genes, whereas PPI is usually provided in a network format. Integrating these two formats is not straightforward. Mainly, two approaches were tried: the matrix-based (MB) method and the network-based (NB) method (Fig. 1).

**Figure 1.**
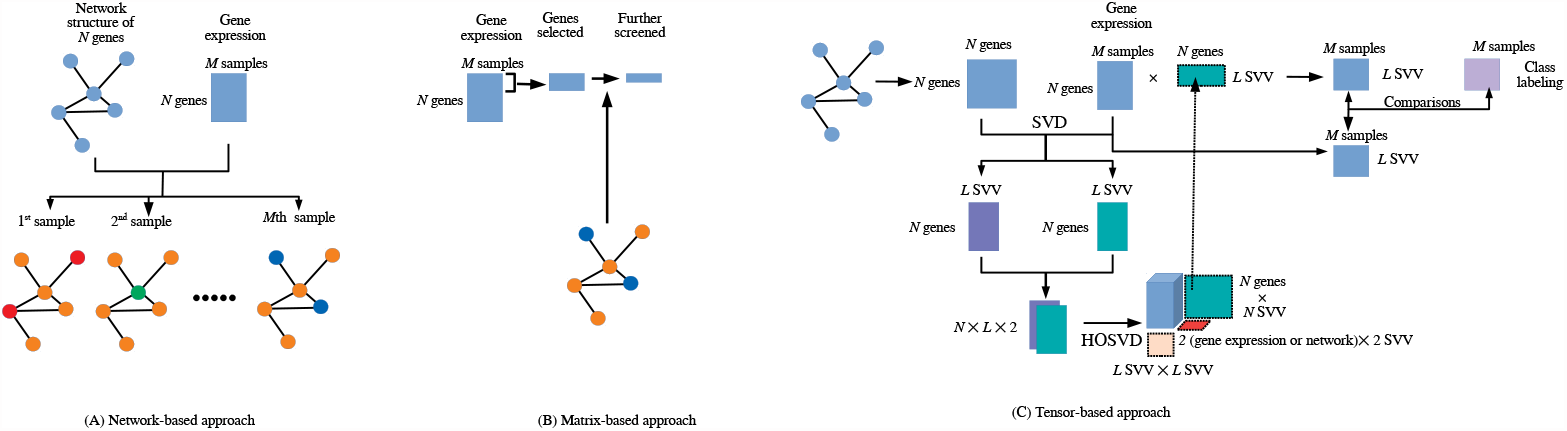
(A) Network-based approach (B) Matrix-based approach (C) Tensor-based approach (this also shows the analysis flow chart in this study)

Typically, in MB methods, differentially expressed genes (DEGs) are identified and PPI is used to validate or filter DEGs [2–4]. However, these kinds of MB methods are rarely recognized as integrated analyses of GE and PPI, since the strategy of further screening DEGs is based on an independent strategy that is common and not restricted to PPI. For example, various enrichment analyses based on previous biological knowledge were often used to screen DEGs. Thus, MB methods are typically unlikely to be regarded as integrated analyses of GE and PPI. In actuality, although [1] reviewed some studies that were regarded as integrated analyses of GE and PPI, most of these studies are not MB but NB methods.

In contrast to MB methods, in NB methods, GE information is mapped onto a network, and modules associated with co-expression genes are selected. These NB approaches integrated with GE are known to improve the performance of simple NB approaches [5–7]. This strategy, where genes embedded in network structures that are further screened based on GE, is more likely to be regarded as integrated analyses of GE and PPI than MB methods, because GE is not the only criterion to further screen genes embedded in a network. Many other criteria, such as the previously mentioned enriched analyses, can be used to further screen genes embedded in a network, and NB methods are not specifically integrated analyses of GE and PPI.

The weak point of both the MB and NB methods is obvious. One of two criteria, GE or PPI, inevitably dominates another. In MB approaches, no non-DEGs are selected because only DEGs are passed to be screened by PPI, whereas in NB approaches, no non-network-connected genes are selected because only genes connected within a network are passed to be screened based on whether they are DEGs. Thus, there are inevitable inequalities between GE and PPI in both approaches.

Some studies attempted to avoid the inequality between GE and PPI. For example, [8] tried to equally weight GE and PPI by introducing the Jaccard similarity index (JDC), which is a simple product between the edge clustering coefficient computed from the PPI network and the Jaccard coefficient computed from the GE similarity. Although JDC was superior to other existing methods and removing the inequality between GE and PPI could improve performance, using JDC is restricted to the time course dataset and this type of approach (i.e., trying to equally weight GE and PPI) has only been rarely investigated.

In this paper, we introduce new, tensor-based (TB) approaches to integrate PPI and GE, assuming no priority between GE and PPI. In the TB approach, PPI expressed in a matrix format is once transformed using singular value decomposition into singular value vectors (SVV), which are later bundled with SVVs computed from GE with SVD to generate a tensor to which TD is applied. The TD-generated SVVs attributed to a gene are further used to generate vectors attributed to samples by projecting GE to TD-generated SVVs attributed to a gene. These obtained vectors attributed to samples are tested if they are coincident with class labels (e.g., patients vs healthy controls) and are selected based on the coincidence. Once vectors attributed to samples are selected, then corresponding SVVs attributed to a gene, to which GE is mapped, is used to select DEGs using previously proposed TD-based unsupervised feature extraction (FE) [9] criterion that was recently improved with optimized standard deviation [10,11] used in Gaussian distribution, which SVVs attributed to genes are supposed to obey in the null hypothesis. Selected genes can be further validated with various enrichment analyses based on previously obtained biological knowledge.

As the scarcity of data hinders the performance of data-driven methods and thereby affecting the biological reliability of selected genes, we proposed a new computational method to integrate GE and PPI and improve the biological significance of obtained results from enrichment analysis as shown from the analysis of cancer data obtained from the GEO database.

## 2. Materials and Methods

In this paper, to express network structure in a matrix format to be integrated with GE, we employed the simplest Network structure representation (NSR) [12], where *n*_*ii*′_ ∈ ℝ^*N*×*N*^ is 1 only when the node *i* and *i*′ are connected with each other and otherwise is zero.

### 2.1. Integrated analysis of matrices and networks with tensor

To convert an NSR into a matrix starting with *n*_*ii*′_, we applied SVD to *n*_*ii*_′ as

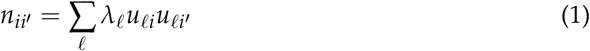

where *λ*_*ℓ*_ is a singular value and *u*_*ℓi*_ ∈ ℝ^*N*×*N*^ are the singular value matrix and the orthogonal matrix.

On the other hand, we apply SVD to *x*_*ij*_ ∈ ℝ^*N*×*M*^, which represents the gene expression of the *i*th gene of the *j*th sample as

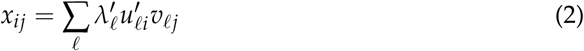

where 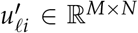 and *υ*_*ℓj*_ ∈ ℝ^*M*×*M*^ (here we assume *M* < *N*). Then, we generate the tensor *x*_*iℓk*_ ∈ ℝ^*N*×*L*×2^ using the first *L*(< *M*) SVVs as:

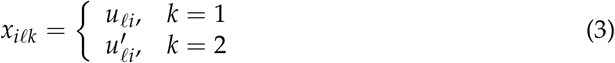

to which higher order singular value decomposition (HOSVD) is then applied, giving:

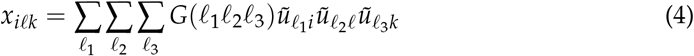

where *G* ∈ ℝ^*N*×*L*×2^ is a core tensor and 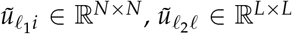, and 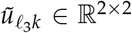 are singular value matrices and are orthogonal matrices. Throughout this article, *L* = 10. Next, we project *x*_*ij*_ onto 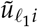 by

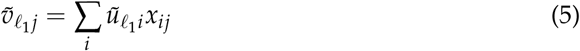

to get vectors attributed to sample, 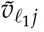.

### 2.2. Comparisons of coincidence with class labels between *υ*_*ℓj*_ and 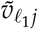

Categorical regression is performed for *υ*_*ℓj*_ and 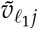 as

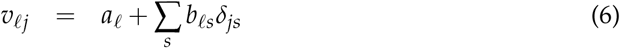

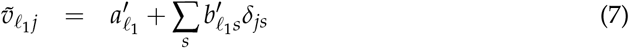

where *δ*_*js*_ is 1 only when *j* belongs to the *s*th class and otherwise is 0, and 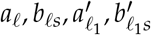 are regression coefficients. *P*-values are computed by the lm function in R. Hereafter, we denote *P*-values computed using Eqs (6) and (7) for the *m*th dataset as 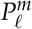 and 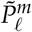, respectively. The obtained *P*-values are corrected by the BH criterion [9] using the p.adjust function in R with the “BH” option.

If there are *M*_0_ data sets (i.e., *m ≤ M*_0_), which is the number of cancer types in this study as shown in the following, then there are *M*_0_*L P*-values (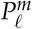 or 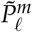), 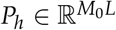which are ranked in ascending order (i.e., if *h* > *h*′ then *P*_*h*_ > *P*_*h*_′). *P*_*h*_s re-ranked from 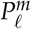 and 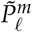 are compared with a paired *t*-test and a paired Wilcoxon test with the alternative hypothesis that the *P*_*h*_ found via SVD (i.e., re-ranked 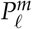) is greater (i.e., less significant) than that given by HOSVD (i.e., re-ranked 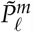).

The coincidence with class labels can be evaluated as follows. *υ*_*ℓj*_ as well as 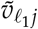 can be regarded as the individual genes’ representative profiles that represent dependence upon *j*. Thus, if *υ*_*ℓj*_ as well as 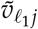have dependence upon class labels, we can regard that individual genes’ profiles have sufficient projection onto those coincident with class labels as well. Moreover, since *υ*_*ℓj*_ or 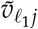 can be computed by projecting *x*_*ij*_ onto 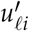or 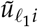, respectively, we can derive the dependence of *x*_*ij*_ upon *j* even only from the dependence upon *i*. In this sense, the valuating coincident of *υ*_*ℓj*_ as well as 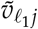 with class labels can be regarded as a measure of inherent coincidence between individual genes’ profiles and class labels as well.

### 2.3. Identification of genes expressed distinctly between class labels and enrichment analysis

First, for each cancer dataset, using one of fives class labels (see below), *ℓ* or *ℓ*_1_ with the smallest 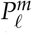 or 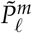 computed by categorical regression of Eqs (6) and (7), respectively, we attributed *P*-values to individual genes (proteins) as follows:

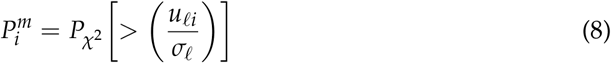

or

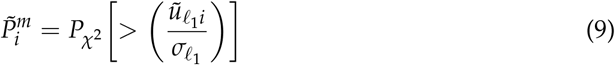

where 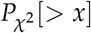 is the cumulative *χ*^2^ distribution when the argument is larger than *x* and *σ*_*ℓ*_ and 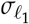 are optimized standard deviations so that 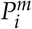 or 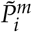 obeys Gaussian, which is the null hypothesis, as much as possible [10,11]. Computed *P*-values are corrected by the BH criterion [9] and *i*s associated with adjusted *P*-values less than 0.01 are selected. Selected genes in each of the 27 cancer types were separately uploaded to Enrichr [13] with the enrichR [14] package. Then, enrichment in KEGG, GO BP, GO CC, and GO MF were retrieved.

Enrichr evaluates enrichment of genes using Fisher’s exact test. Suppose *G*_1_ is a set of genes uploaded (i.e., genes selected by our method) and *G*_2_ is a set of genes with known function (e.g., genes that belong to a specific KEGG pathway). The overlap between *G*_1_ and *G*_2_, *G*_1_ ∩ *G*_2_, is evaluated by the comparison with that by chance. If *G*_1_ ∩ *G*_2_ is much larger than that by chance and the probability of occurrence by chance is small, *G*_1_ is regarded to be associated with the function associated with *G*_2_. In this study, we selected biological terms enriched if associated adjusted *P*-values given by Enrichr is less than 0.05.

### 2.4. PPI dataset

We have employed the following two PPI datasets for comparisons.

#### 2.4.1. Stanford PPI dataset

The PPI dataset, PP-Pathways_ppi.csv.gz, was retrieved from the human protein-protein interaction network [15], which includes 342,353 pairs for 21,557 proteins. After excluding self-pairs (i.e., self-dimers), these data were formatted as a *n*_*ii*_ ∈ ℝ^*N*×*N*^ where *N* = 21557. Then, there were 16,774 *i*s, which is common with the *i*s in the TCGA gene expression profiles (see below). Since these 342,353 pairs represent only one of *n*_*ij*_ and *n*_*ji*_, (i.e., pairs whose order is reversed are not included), when *n*_*ij*_ ≠ *n*_*ji*_, *n*_*ij*_ = *n*_*ji*_ = 1 is required.

#### 2.4.2. BIOGRID PPI dataset

The PPI dataset, BIOGRID-MV-Physical-4.4.221.tab2.zip, which is supposed to represent physical PPIs, was retrieved from BIOGRID [16], which includes 437,679 pairs for 27,978 proteins. These data were also formatted as a *n*_*ii*_′ ∈ ℝ^*N*×*N*^ where *N* = 27978. Then, there were 11,294 *i*s, which is common with the *i*s in the TCGA gene expression profiles (see below). Since these 437,679 pairs represent only one of *n*_*ij*_ and *n*_*ji*_, (i.e., pairs whose order is reversed are not included), when *n*_*ij*_ ≠ *n*_*ji*_, *n*_*ij*_ = *n*_*ji*_ is required. In the BIOGRID dataset, *n*_*ij*_ is not always taken to be 1 when protein pairs interact with each other, but the number of occurrences in the BIOGRID PPI datasets. Thus, *n*_*ij*_ can be larger than 1 in contrast to the Stanford PPI. Thus, *n*_*ij*_ in the BIOGRID PPI represents not only if pairs of proteins interact with each other, but also the strength of the interaction.

### 2.5. Gene expression profiles

TCGA gene expression profiles are used as *x*_*ij*_. The RTCGA dataset [17] was used for this purpose. RTCGA.rnaseq [18] was used as a gene expression profile. It includes 27 cancer datasets with various sample sizes (*j*s) ranging from a few tens to a few hundred, as well as 20,532 genes (*i*s) whose expression profiles are available. The cancers considered are ACC, BLCA, BRCA, CESC, COAD, ESCA, GBM, HNSC, KICH, KIRC, KIRP, LGG, LIHC, LUAD, LUSC, OV, PAAD, PCPG, PRAD, READ, SARC, SKCM, STAD, TGCT, THCA, UCEC, and UCS. The class labels considered, retrieved from RTCGA.clinical [19], are patient.vital_status, patient.stage_event.pathologic_stage, patient.stage_event.tnm_categories. pathologic_categories. pathologic_m, patient.stage_event.tnm_categories.pathologic_categor pathologic_n, and patient.stage_event.tnm_categories. pathologic_categories. pathologic_t. In order to avoid complexity, in the following, we employ shortened label class names as follows: “vital_status,” “pathologic_stage,” “pathologic_m,” “pathologic_n,” and “pathologic_t.” All 27 cancer datasets are associated with “vital_status” labels, “pathologic_stage” and “pathologic_m” are associated with only 18 datasets, and “pathologic_t” and “pathologic_n” are associated with 20 datasets (Table 1).

**Table 1.**
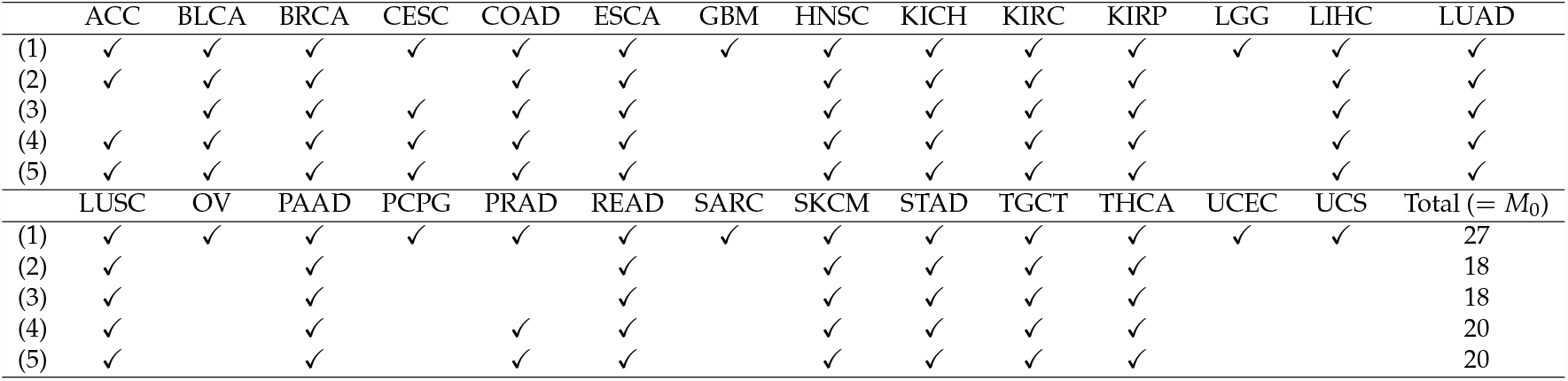
Availability of class labels for cancer datasets. (1) “patient.vital_status,” (2) “pathologic_stage,” (3)“pathologic_m,” (4) “pathologic_t,” and (5) “pathologic_n”.

16,774 *i*s (for stanford) and 11,294 (for BIOGRID) that are common with *i*s in PPI (see above) are used.

## 3. Results

### 3.1. Identification of sample vectors coincident with labels

The first evaluation of the proposed TB method is a comparison of the significance of coincidence between labels and vectors attributed to samples with and without a consideration of PPI (i.e., comparisons between Eqs (6) and (7)). If Eq. (7) can provide more significance than Eq. (6), an integrated analysis of PPI and GE can improve the performance, because PPI itself is unlikely to include class label information, which is supposed to be specific to individual cancer types. The reason why we employed these class labels is simply because they are widely common for majority of cancer types in TCGA. Since they are patients class labels, they might not be directly related to some specific biological concepts.

#### 3.1.1. Stanford PPI

In this section, we each evaluate class label.

##### “vital_status”

First, we considered the label “vital_status”, which has two levels, “dead” and “alive.” Figure 2 (the left panel, (1)) represents the logarithmic *P*-values computed by applying a *t*-test (2.585 × 10^*−*12^) and a Wilcoxon test (2.144 × 10^*−*8^) to ascending ordered log_10_ *P*_*h*_ computed from *υ*_*ℓj*_ with Eq. (6) and 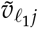with Eq. (7), respectively, whose scatter plot is shown in Fig. 3 (the left panel). Since the number of vectors attributed to samples associated with adjusted *P*-values less than 0.05 for HOSVD is larger than those for SVD, the integrated analysis clearly improves the coincidence between the class label “vital_status” and vectors attributed to samples.

**Figure 2.**
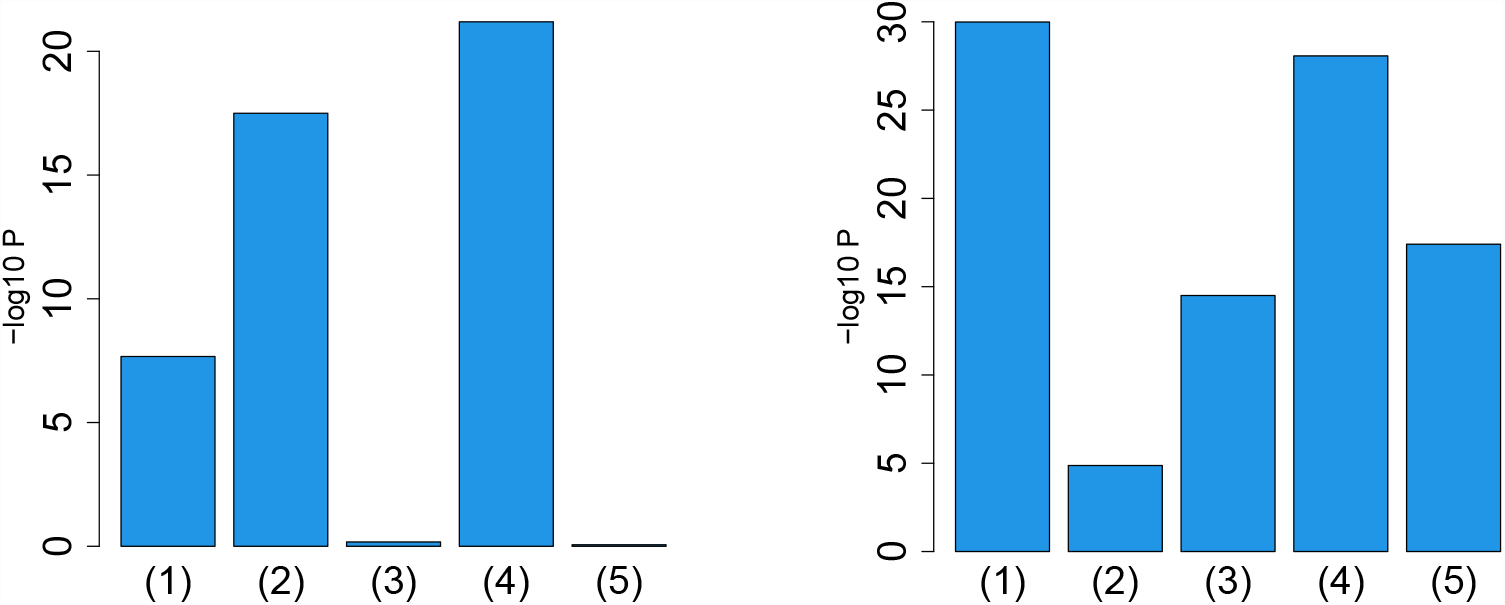
Barplot of *P*-values computed by a Wilcoxon test to evaluate the difference in ascending ordered *P*_*h*_ between SVD (Eq. (6)) and HOSVD (Eq. (7)) when Stanford PPI (left) or BIOGRID PPI (right) was used. (1) “vital_status”, (2) “pathologic_stage”, (3) “pathologic_m” (4) “pathologic_t” (5) “pathologic_n”. Numerical values of bar plots are listed in Table S1

**Figure 3.**
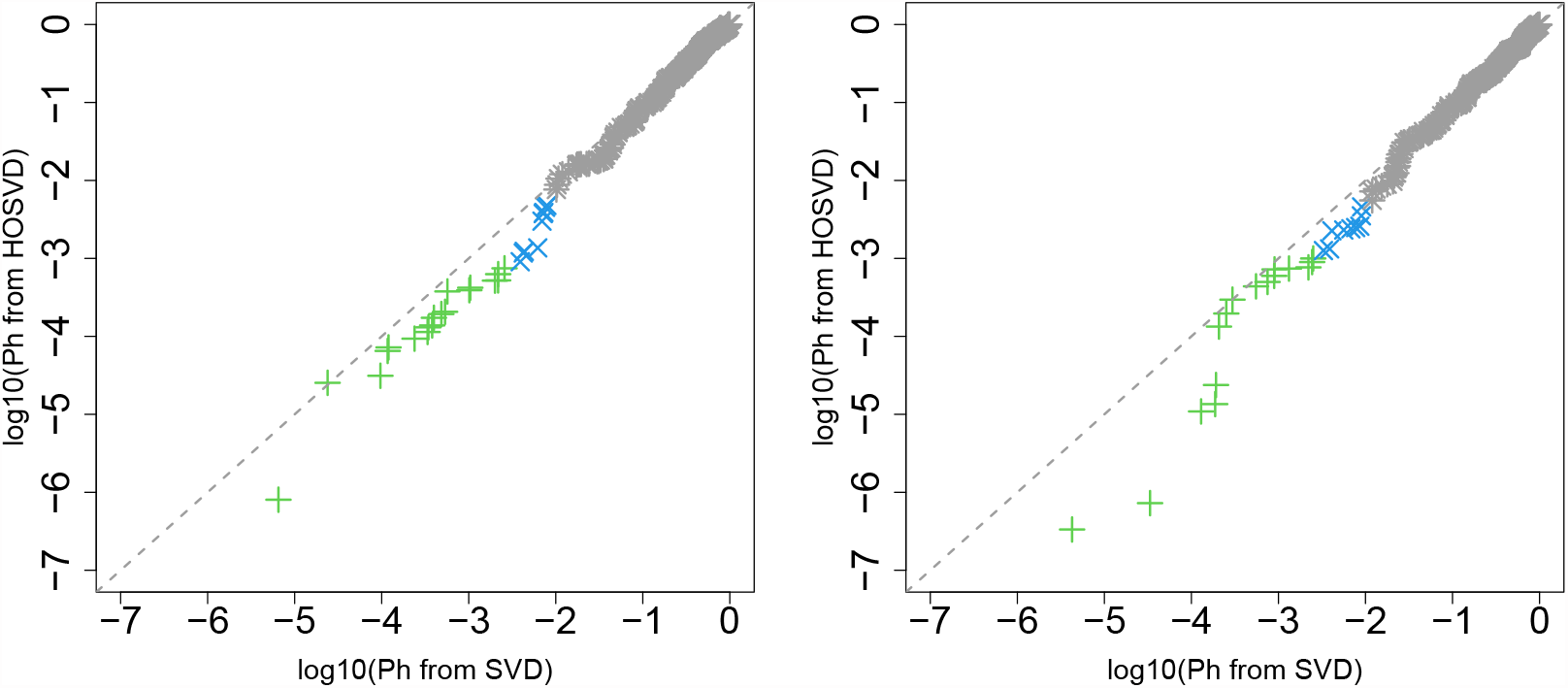
Scatter plot (logarithmic scale) of ascending ordered *P*_*h*_ computed from *υ*_*ℓj*_ (horizontal axis) and 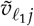 (vertical axis) for vital_status. Green crosses are those associated with adjusted *P*-values less than 0.05 for both axes and blue crosses are those associated with adjusted *P*-values less than 0.05 for the vertical axis alone. Grey asterisks represent all other situations. Left: Stanford PPI, right: BIOGDID PPI

##### “pathologic_stage”

Next, we considered the label “pathologic_stage.” Figure 2 (the left panel, (2)) represents the logarithmic *P*-values computed by applying a *t*-test (1.791×10^*−*5^) and a Wilcoxon test (3.178×10^*−*18^) to ascending ordered log_10_ *P*_*h*_ computed from *υ*_*ℓj*_ with Eq. (6) and 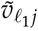 with Eq. (7), respectively, whose scatter plot is shown in Fig. 4 (the left panel). Again, *P*_*h*_ with integrated analysis of GE and PPI is significantly lower than that without consideration of PPI. Thus, the improvement observed in the label “vital_status” is unlikely accidental.

**Figure 4.**
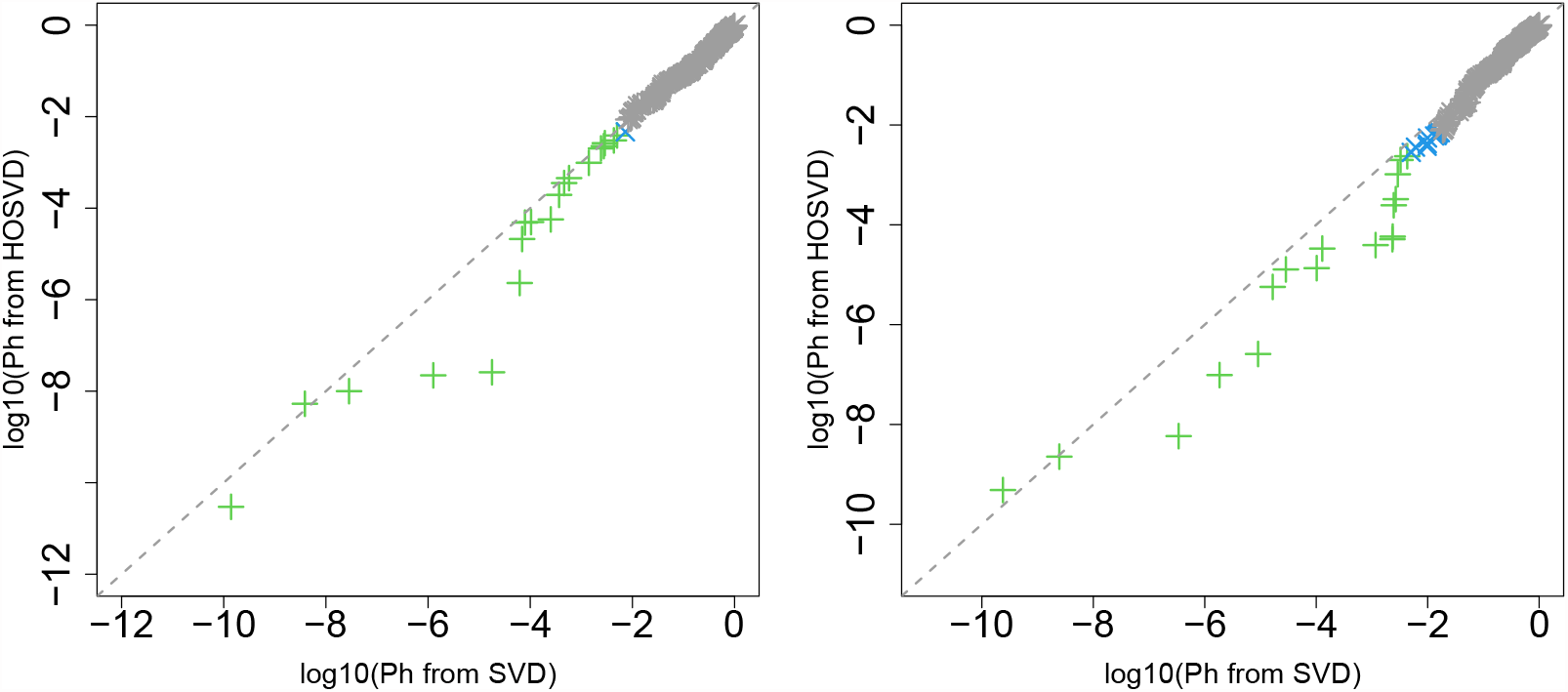
Scatter plot in logarithmic scale of ascending ordered *P*_*h*_ computed from *υ*_*ℓj*_ (horizontal axis) and 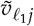 (vertical axis) for “pathologic_stage.” Green crosses are those associated with adjusted *P*-values less than 0.05 for both axes and blue crosses are those associated with adjusted *P*-values less than 0.05 only for vertical axis. Grey asterisks represent all other situations. Left: Stanford PPI, right: BIOGRID PPI

##### “pathologic_m”

Next, we considered the label “pathologic_m.” Figure 2 (the left panel, (3)) represents the logarithmic *P*-values computed by applying a *t*-test (0.6714) and a Wilcoxon test (0.6685) to ascending ordered log_10_ *P*_*h*_ computed from *υ*_*ℓj*_ with Eq. (6) and 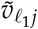 with Eq. (7), respectively, whose scatter plot is shown in Fig. 5 (the left panel). Since consideration of PPI does not improve the coincidence between class labels and vectors attributed to samples in this case, integrated analysis of PPI and GE does not always improve the coincidence (this will be discussed further).

**Figure 5.**
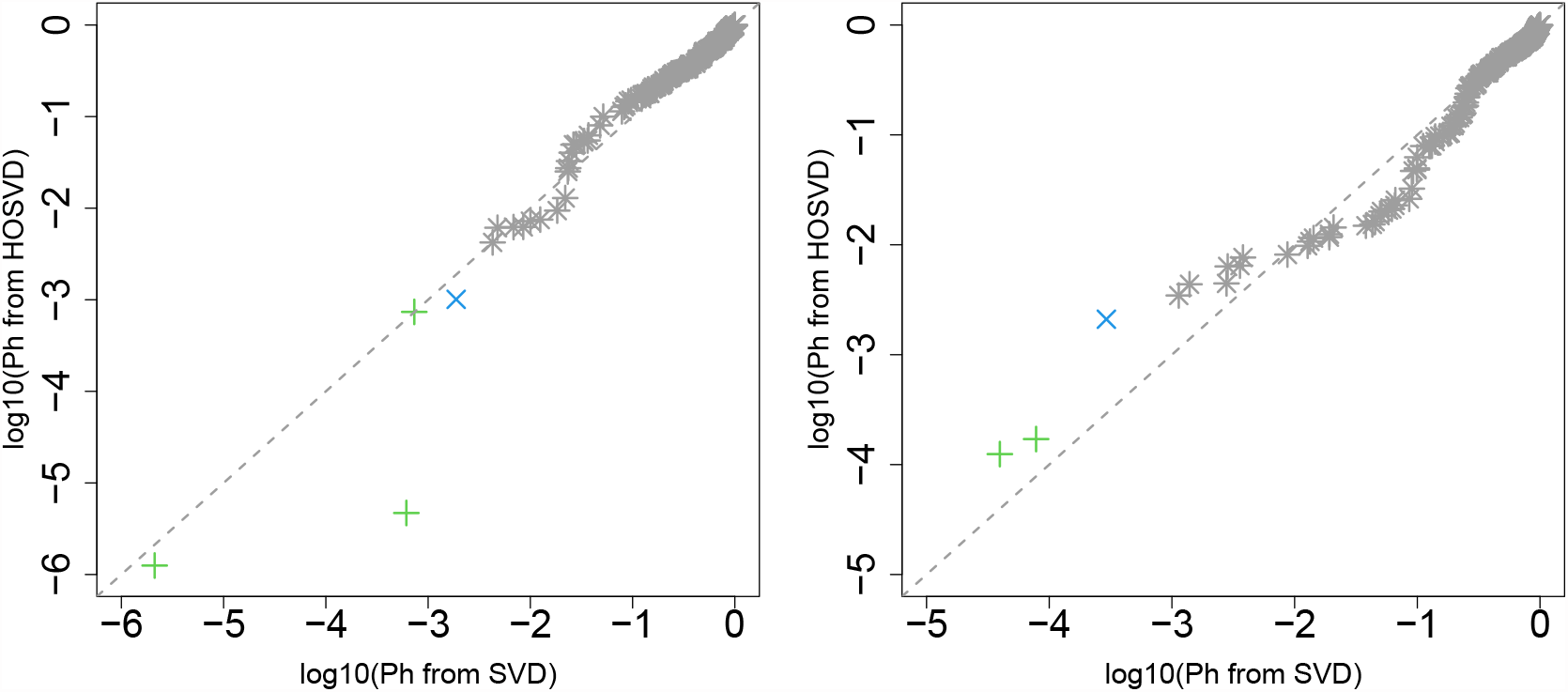
Scatter plot (logarithmic scale) of ascending ordered *P*_*h*_ computed from *υ*_*ℓj*_ (horizontal axis) and 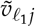 (vertical axis) for “pathologic_m.” Green crosses are associated with adjusted *P*-values less than 0.05 for both axes, and blue crosses are associated with adjusted *P*-values less than 0.05 for the vertical axis alone. Grey asterisks represent all other situations. Left: Stanford PPI, right: BIOGRID PPI.

##### “pathologic_t”

Next, we considered the label “pathologic_t.” Figure 2 (the left panel, (4)) represents the logarithmic *P*-values computed by applying a *t*-test (2.591×10^*−*3^) and a Wilcoxon test (6.430×10^*−*22^) to ascending ordered log_10_ *P*_*h*_ computed from *υ*_*ℓj*_ with Eq. (6) and 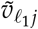 with Eq. (7), respectively, whose scatter plot is shown in Fig. 6 (the left panel). Although the log *P*_*h*_ does not appear significantly distinct between SVD and HOSVD (Fig. 6, the left panel), since the *P*-values are small enough (Fig. 2, the left panel, (4)), integrated analysis of PPI and GE could improve the coincidence between vectors attributed to samples with class labels.

**Figure 6.**
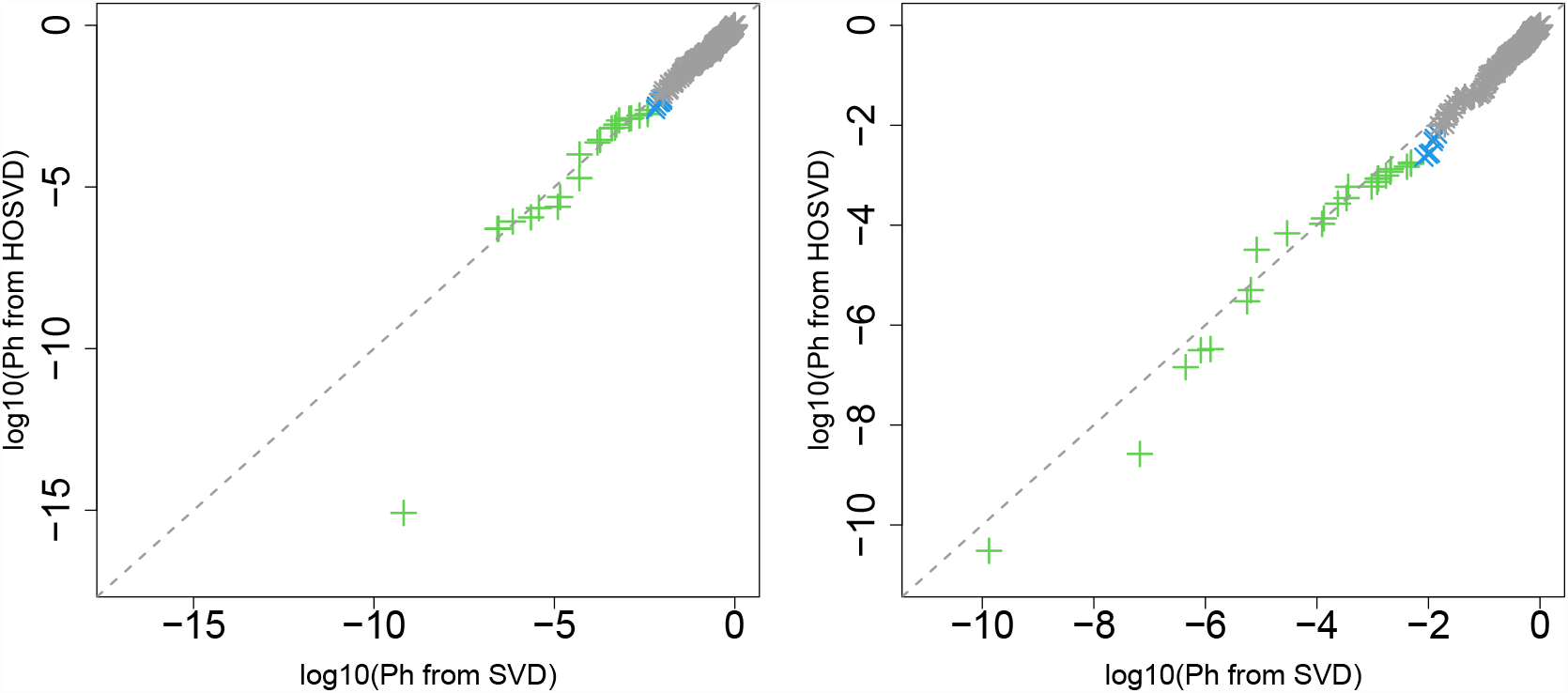
Scatter plot (logarithmic scale) of ascending ordered *P*_*h*_ computed from *υ*_*ℓj*_ (horizontal axis) and 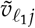 (vertical axis) for “pathologic_t.” Green crosses are associated with adjusted *P*-values less than 0.05 for both axes, and blue crosses are associated with adjusted *P*-values less than 0.05 for the vertical axis alone. Grey asterisks represent all other situations. Left: Stanford PPI, right: BIOGRID PPI

##### “pathologic_n”

Next, we considered the label “pathologic_n.” Figure 2 (the left panel, (5)) represents the logarithmic *P*-values computed by applying a *t*-test (9.111 × 10^*−*3^) and a Wilcoxon test (0.8667) to ascending ordered log_10_ *P*_*h*_ computed from *υ*_*ℓj*_ with Eq. (6) and 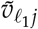 with Eq. (7), respectively, whose scatter plot is shown in Fig. 7 (the left panel). Since most vectors attributed to samples associated with adjusted *P*_*h*_ less than 0.05 for SVD are associated with lower *P*_*h*_ for HOSVD, integrated analysis of GE and PPI surely improved the coincidence between vectors attributed to samples and class labels.

**Figure 7.**
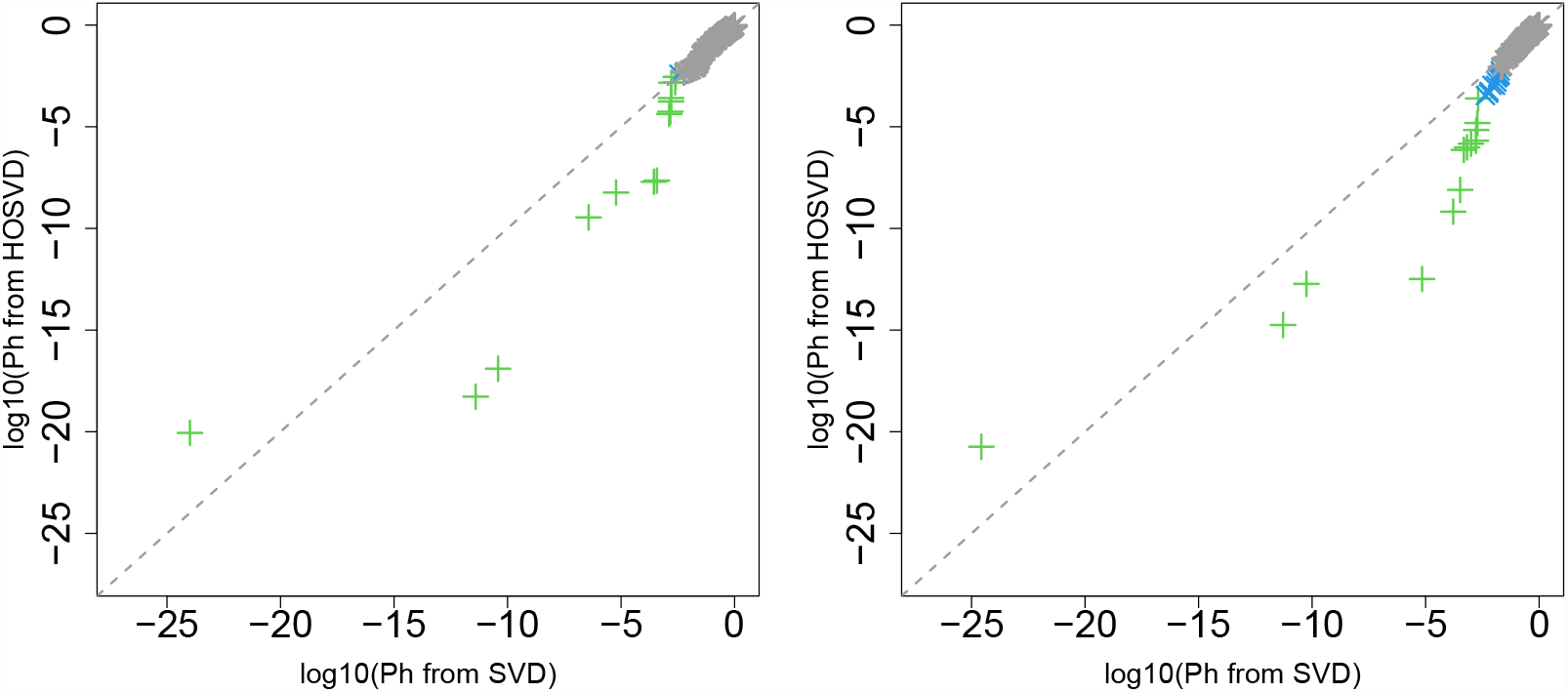
Scatter plot (logarithmic scale) of ascending ordered *P*_*h*_ computed from *υ*_*ℓj*_ (horizontal axis) and 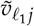 (vertical axis) for “pathologic_n.” Green crosses are associated with adjusted *P*-values less than 0.05 for both axes, and blue crosses are associated with adjusted *P*-values less than 0.05 for the vertical axis alone. Grey asterisks represent all other situations. Left: Stanford PPI, right: BIOGRID PPI

#### 3.1.2. BIOGRID PPI

Although integrated analysis of Stanford PPI and GE surely improved the coincidence between vectors attributed to samples and class labels for four out of five class labels (Fig. 2), we were not certain if one class label, “pathologic_m”, without improved coincidence is because of PPI or because of the class label itself. Furthermore, we were not certain whether coincidence increased or decreased when we considered other PPIs. To examine these questions, we tested another PPI taken from BIOGRID. In the following section, we evaluate each class label.

##### “vital_status”

First, we considered the label “vital_status,” which has two levels, “dead” and “alive.” Figure 2 (the right panel, (1)) represents the logarithmic *P*-values computed by applying a *t*-test (1.696 × 10^*−*12^) and a Wilcoxon test (1.027451 × 10^*−*30^) to ascending ordered log_10_ *P*_*h*_ computed from *υ*_*ℓj*_ with Eq. (6) and 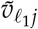 with Eq. (7), respectively, whose scatter plot is shown in Fig. 3 (the right panel). Compared to the improvement when Stanford PPI is used (left panel in Fig. 3), coincidence improvement increased when PPI is considered. This suggests that which PPI is used greatly affects the improvement when integrated analysis of GE and PPI is performed.

##### “pathologic_stage”

Next, we considered the label “pathologic_stage.” Figure 2 (the right panel, (2)) represents the logarithmic *P*-values computed by applying *t*-test (3.644 × 10^*−*7^) and a Wilcoxon test (1.34 × 10^*−*5^) to ascending ordered log_10_ *P*_*h*_ computed from *υ*_*ℓj*_ with Eq. (6) and 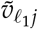 with Eq. (7), respectively, whose scatter plot is shown in Fig. 4 (the right panel). Compared to Fig. 4, coincidence improvement slightly decreased when PPI is considered. Although the direction of alteration is opposite to the “vital_status,” this suggests again that which PPI is used greatly affects the improvement when integrated analysis of GE and PPI is performed.

##### “pathologic_m”

Next, we considered the label “pathologic_m.” Figure 2 (the right panel, (3)) represents the logarithmic *P*-values computed by applying a *t*-test (9.49 × 10^*−*6^) and a Wilcoxon test (3.16 × 10^*−*15^) to ascending ordered log_10_ *P*_*h*_ computed from *υ*_*ℓj*_ with Eq. (6) and 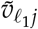 with Eq. (7), respectively, whose scatter plot is shown in Fig. 5 (the right panel). Although *P*-values computed by the *t*-test and the Wilcoxon test are small enough to be significant, because no *P*_*h*_s associated with significant adjusted *P*-values (the green and blue crosses in the right panels in Fig. 5) decreased because of integrated analysis of PPI and GE, it is not regarded as improvement. Thus, the failure of “pathologic_m” when Stanford PPI was used (left panel in Fig. 5) is likely because of the class label itself and not because of PPI.

##### “pathologic_t”

Next, we considered the label “pathologic_t.” Figure 2 (the right panel, (4)) represents the logarithmic *P*-values computed by applying a *t*-test (9.99 × 10^*−*16^) and a Wilcoxon test (8.534979 × 10^*−*29^) to ascending ordered log_10_ *P*_*h*_ computed from *υ*_*ℓj*_ with Eq. (6) and 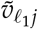 with Eq. (7), respectively, whose scatter plot is shown in Fig. 6 (the right panel). Although two panels in Fig. 6 do not look distinct, since *P*-values computed by the *t*-test and the Wilcoxon test (Fig. 2 (4) in the right panel) are much smaller than those when Stanford PPI was employed (Fig. 2 (4) in the left panel), integrated analysis of PPI and GE could improve the coincidence between vectors attributed to samples with class labels.

##### “pathologic_n”

Next, we considered the label “pathologic_n.” Figure 2 (the right panel, (5)) represents the logarithmic *P*-values computed by applying a *t*-test (3.437 × 10^*−*5^) and a Wilcoxon test (3.930971 × 10^*−*18^) to ascending ordered log_10_ *P*_*h*_ computed from *υ*_*ℓj*_ with Eq. (6) and 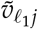 with Eq. (7), respectively, whose scatter plot is shown in right panel of Fig. 7 (the right panel). Because the right panel of Fig. 7 is somewhat improved, compared to the left panel of Fig. 7, the employment of BIOGRID PPI could improve the performance with Stanford PPI.

### 3.2. Identification of DEGs and enrichment analysis

In general, integrated analysis of PPI and GE could improve the coincidence between vectors attributed to samples and class labels. Nevertheless, it is still unclear whether the improved coincidence between vectors attributed to samples and class labels is useful. To address this problem, we have performed enrichment analysis of DEGs.

#### 3.2.1. Stanford PPI

Figure 8 shows the summation of the number of biological terms enriched over 27 cancer classes for four categories (KEGG, GO BP, GO CC, and GO MF). A more detailed cancer cell barplot is available as supplementary material. It is obvious that integrated analysis of PPI and GE increases the number of enriched biological terms, no matter which category is considered.

**Figure 8.**
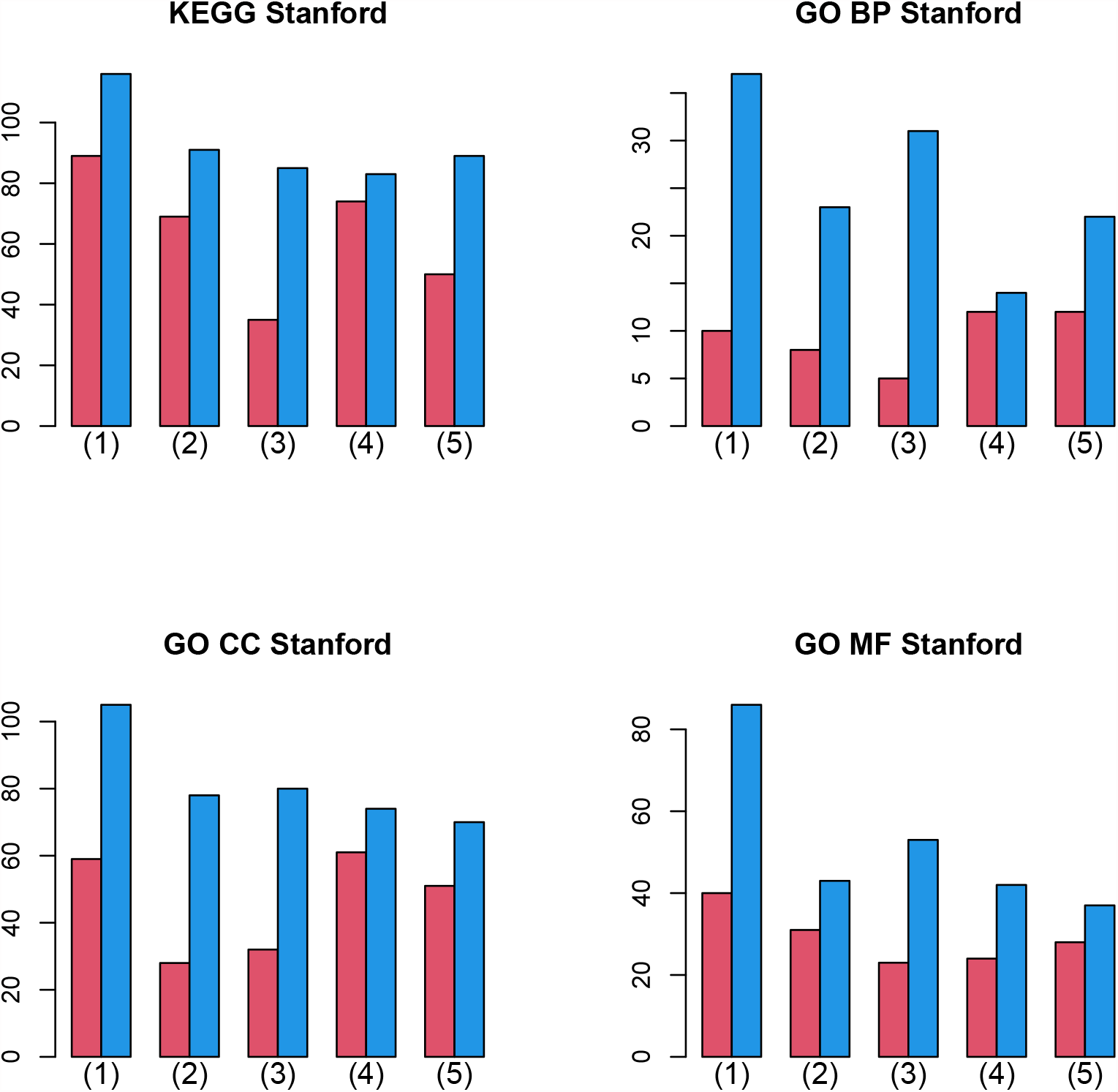
Barplot of the number of enriched biological terms summed over 27 cancers when Stanford PPI was used. (1) “vital_status,” (2) “pathologic_stage,” (3) “pathologic_m,” (4) “pathologic_t,” and (5) “pathologic_n.” Red: without integration of PPI, blue: with integration of PPI.

#### 3.2.2. BIOGRID PPI

Figure 9 shows the summation of the number of biological terms enriched over 27 cancer classes for four categories (KEGG, GO BP, GO CC, and GO MF). A more detailed cancer cell barplot is available as supplementary material. It is obvious that integrated analysis of PPI and GE increases the number of enriched biological terms excluding a few cases, although the amount of the increase is less than in Fig. 8.

**Figure 9.**
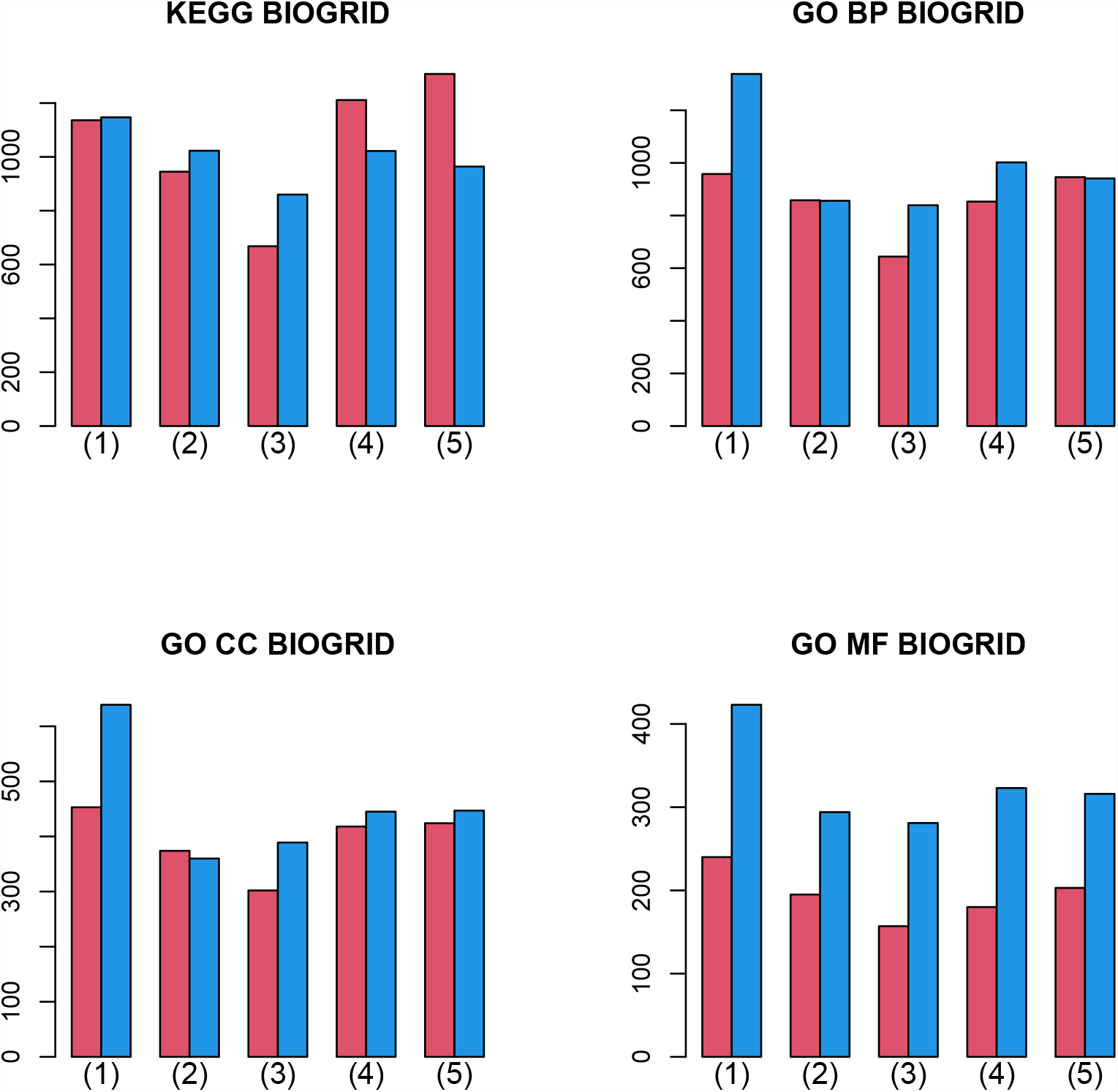
Barplot of the number of enriched biological terms summed over 27 cancers when BIOGRID PPI was used. (1) “vital_status,” (2) “pathologic_stage,” (3) “pathologic_m,” (4) “pathologic_t,” and (5) “pathologic_n.” Red: without integration of PPI, blue: with integration of PPI.

In conclusion, the integrated analysis of PPI and GE increases not only the coincidence between vectors attributed to samples and class labels, but also the biological reliability of selected genes.

## 4. Discussion

Integrated analysis of PPI and GE is unlikely to improve coincidence with class labels, because there are many class labels which are distinct from one another, whereas the PPI network does not vary depending on class labels. GE also does not change its values depending on class labels, but because samples are associated with class labels, it is not surprising that GE is coincident with class labels to some extent. Nevertheless, why can PPI that does not have any direct relation to class labels improve coincidence with class labels?

To clarify this point, we tried a simpler integration between PPI and GE. We computed sample vectors directly from PPI, not passing through TD. Mathematically, vectors attributed to samples can be computed directly from PPI as

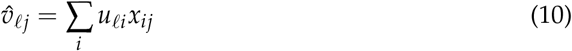

where *u*_*ℓi*_ was computed from PPI, *n*_*ii*_, with Eq. (1). Since *x*_*ij*_ is GE and *u*_*ℓi*_ comes from PPI, it is a type of integrated analysis of GE and PPI. Then, coincidence with class labels is evaluated using categorical regression as

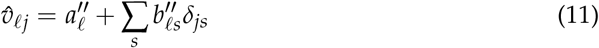

which just has been done in the above analysis.

Next, we investigated the correlation between 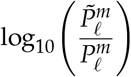 and 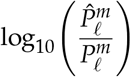 over 27 cancers, where 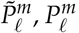, and 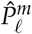 are *P*-values computed from Eqs (7), (6), and (11) for *m*th cancer type. This correlation evaluated whether the coincidence improvement between vectors attributed to samples and class labels by integrated analysis with TD, 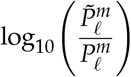, correlates with the coincidence improvement of the integrated analysis without TD evaluated by Eq. (11) from coincidence in GE only evaluated by Eq. (6), 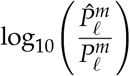. Interestingly, regardless of class labels and PPI, there are significant correlations (Fig. 10). This means that improved coincidence between vectors attributed to samples and class labels caused by integrated analysis of PPI and GE with TD can be observed even in a simpler integration of PPI and GE (Eq. (10)) to some extent, and this is why TB integrated analysis of PPI and GE can improve the coincidence between vectors attributed to samples and class labels.

**Figure 10.**
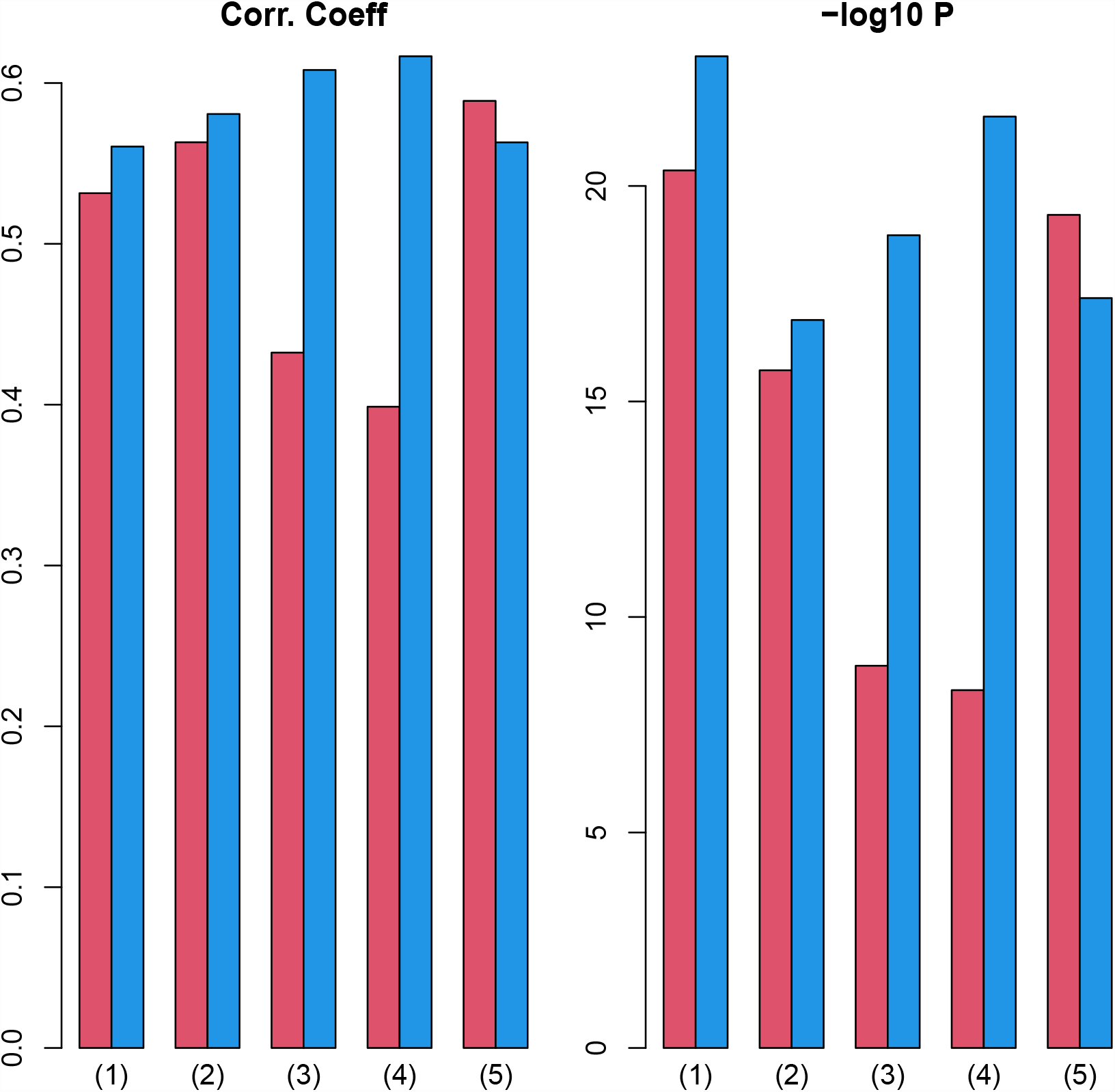
Left: Pearson correlation coefficient between 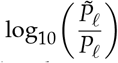and 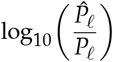over 27 cancers. Right: associated *P*-values (logarithmic scale) (1) “vital_status,” (2) “pathologic_stage,” (3) “pathologic_m,” (4) “pathologic_t,” and (5) “pathologic_n.” Red: Stanford PPI, blue: BIOGRID PPI. Numerical values of bar plots are listed in Table S2.

Since one might wonder if simpler a integration of PPI and GE (Eq. (10)) is powerful enough to improve the coincidence between vectors attributed to samples and class labels even without TD, we evaluated (Fig. 11) the improvement of the coincidence using a simpler integration of PPI and GE, as in Figs 2. Since it is obvious that simpler integration of PPI and GE with Eq. (10) is less likely to improve the coincidence between vectors attributed to samples and class labels than TB integration, TB integration is required to improve the coincidence between vectors attributed to samples and class labels.

**Figure 11.**
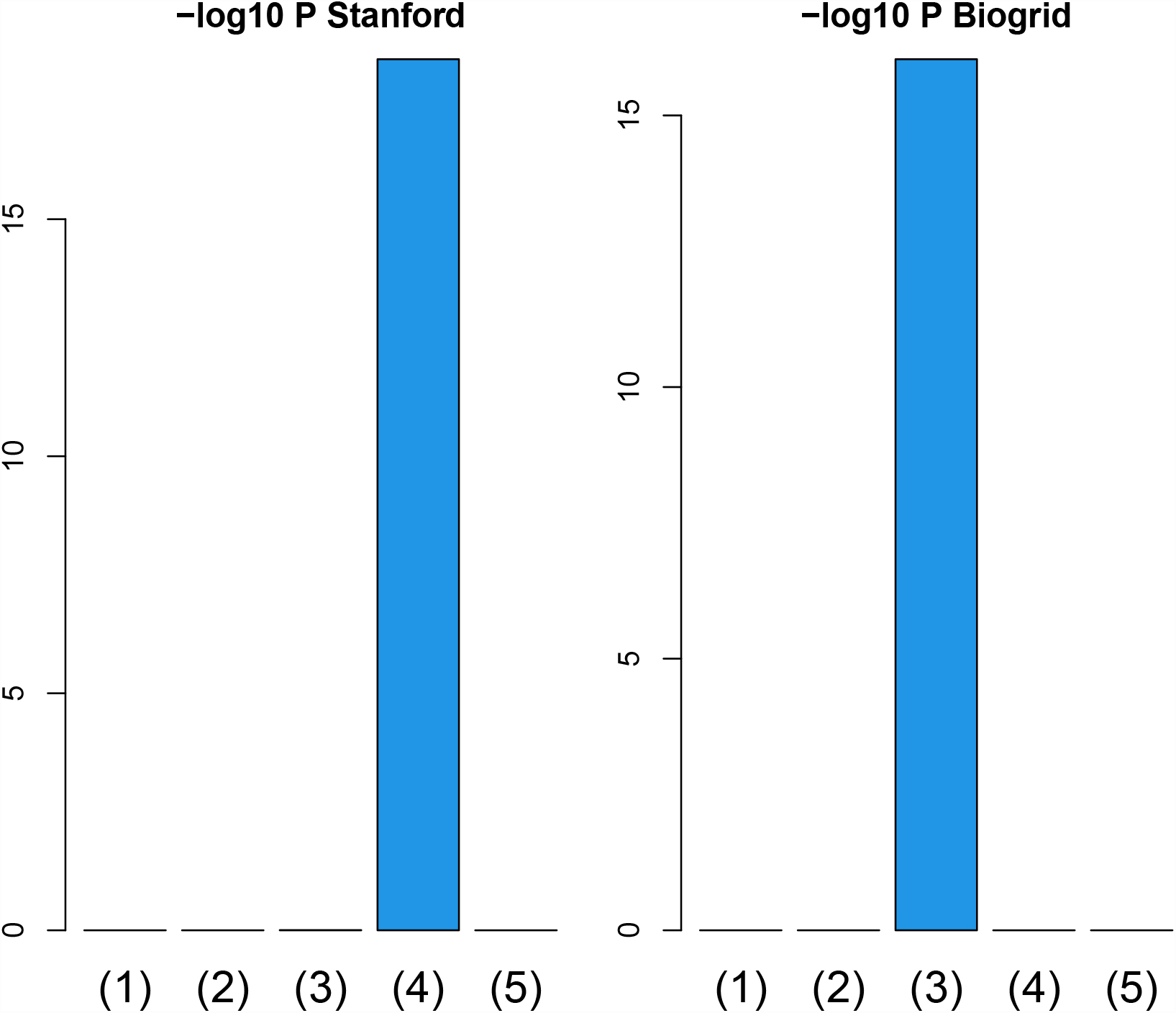
Barplot of *P*-values computed by a Wilcoxon test to evaluate the difference in ascending ordered *P*_*h*_ between SVD (Eq. (6)) and simpler integration (Eq. (11)). (1) “vital_status,” (2) “pathologic_stage,” (3) “pathologic_m,” (4) “pathologic_t,” and (5) “pathologic_n.” Left: Stanford PPI, right: BIOGRID PPI. Numerical values of bar plots are listed in Table S3.

Considering the distinct performances of Stanford PPI (left panel in Fig. 2 and Fig. 8), where *n*_*ij*_ takes only 1 or 0 dependent upon whether pairs of proteins interact or not and BIOGRID PPI (right panel in Fig. 2 and Fig. 9), where *n*_*ij*_ can take larger values than 1 to represent the strength of interaction, it is likely better to consider not only if the interaction exists between pairs of proteins but also how strong the interaction between pairs of protein is. This finding might be able to help us to consider the integration of PPI and GE in the future.

We have also noted that enrichment of SVD based gene selection (i.e., without integration of PPI) in BIOGRID (the blue bars in Fig. 9) is better than in Stanford PPI (the blue bars in Fig. 8). This should not happen, since nothing can change between Stanford PPI and BIOGRID PPI when no integrated analyses were not performed. Nevertheless, PPI can affect outcomes even before integration, since genes are screened based on whether they are also included in PPI. Because Stanford PPI and BIOGRID PPI differ, the other genes considered also differ. It turned out that this gene restriction largely affected the enrichment analysis. In this sense, integrated analysis can affect the outcome, but so does restricting the genes in considering overlaps with PPI.

To see if integrated analysis correctly identifies biologically reasonable genes, we investigated the frequency of selected terms in KEGG pathway for BIOGRID data (Table 2). Pathways in Table 2 are related to cancers. “Pathways in cancer” is directly related to cancer. “JAK-STAT signaling pathway” is related to cancers [20]. As for “Cytokine-cytokine receptor interaction”, Cytokine signaling is known to be related to cancers [21]. “PI3K-Akt signaling pathway” is known to be dysregulated almost in all human cancers [22]. As for “C-type lectin receptor signaling pathway”, C-Type lectin receptors are related to cancer immunity [23]. In addition to these, increased colon cancer risk was observed after severe Salmonella infection [24], cancers are associated with less apotosis [25], non-alcoholic fatty liver disease increases the cancer risk [26], influenza is associated with worse in-hospital clinical outcomes among hospitalised patients with malignancy [27]. Thus most of the frequently selected pathways are related to cancers.

**Table 2.** Ten top most frequently selected pathways in KEGG pathway for BIOGRID data when PPI and GE are integrated. The numbers indicate the frequency of selections among 27 cancer types. (1) “vital_status,” (2) “pathologic_stage,” (3) “pathologic_m,” (4) “pathologic_t,” and (5) “pathologic_n.”

| Pathway | (1) | (2) | (3) | (4) | (5) |
| --- | --- | --- | --- | --- | --- |
| Salmonella infection | 25 | 18 | 17 | 20 | 20 |
| JAK-STAT signaling pathway | 23 | 18 | 17 | 19 | 20 |
| Cytokine-cytokine receptor interaction | 23 | 17 | 17 | 19 | 20 |
| Influenza A | 22 | 17 | 16 | 18 | 19 |
| Pathways in cancer | 22 | 17 | 14 | 19 | 19 |
| Apoptosis | 20 | 18 | 14 | 18 | 16 |
| Ribosome | 20 | 15 | 16 | 16 | 15 |
| Non-alcoholic fatty liver disease | 18 | 16 | 14 | 18 | 16 |
| PI3K-Akt signaling pathway | 16 | 17 | 15 | 18 | 16 |
| C-type lectin receptor signaling pathway | 18 | 15 | 13 | 17 | 16 |

Table 2 included many biological pathways other than cancer specific ones. To see more cancer specific results, we restrict to “HALLMARK cancer gene sets” and repeated the above procedure for the integration with BIOGRID. (Table 3). Although the number of frequency of selection decreased from Table 2 possibly because of more specific (strict) evaluation of the relationship to cancers, Table 3 still includes substantial number of frequency of selection. Moreover, the number of selected gene sets are more in the integrated analysis (denoted as HOSVD) than without integrated analysis (denoted as SVD). Therefore, even if we consider more cancer specific features, the integrated analysis of PPI and GE has substantial number of identtification and more numbers than that without integration

**Table 3.** Enriched gene sets in “HALLMARK cancer gene sets” with and without the integration using BIOGRID. The numbers indicate the frequency of selections among 27 cancer types. (1) “vital_status,” (2) “pathologic_stage,” (3) “pathologic_m,” (4) “pathologic_t,” and (5) “pathologic_n.”

| Pathway | (1) | (2) | (3) | (4) | (5) |
| --- | --- | --- | --- | --- | --- |
| HOSVD (with integration) |  |  |  |  |  |
| HALLMARK_MYC_TARGETS_V1 | 10 | 4 | 5 | 6 | 6 |
| HALLMARK_FATTY_ACID_METABOLISM | 6 | 4 | 6 | 5 | 4 |
| HALLMARK_OXIDATIVE_PHOSPHORYLATION | 3 | 1 | 1 | 2 | 2 |
| HALLMARK_APOPTOSIS | 3 | 1 | 2 | — | 2 |
| HALLMARK_PEROXISOME | 2 | 2 | — | 1 | — |
| HALLMARK_IL6_JAK_STAT3_SIGNALING | 1 | — | 1 | — | — |
| HALLMARK_MYC_TARGETS_V2 | — | — | 1 | — | 1 |
| HALLMARK_ALLOGRAFT_REJECTION | 1 | — | — | — | — |
| HALLMARK_APICAL_SURFACE | 1 | — | — | — | — |
| SVD (without integration) |  |  |  |  |  |
| HALLMARK_APOPTOSIS | 11 | 5 | 7 | 3 | 3 |
| HALLMARK_FATTY_ACID_METABOLISM | 6 | 3 | 4 | 3 | 3 |
| HALLMARK_PEROXISOME | 2 | 2 | — | 3 | 3 |
| HALLMARK_MYC_TARGETS_V1 | 1 | 2 | 2 | 1 | 1 |

## 5. Conclusions

We have proposed integrated analysis of PPI and GE with TD that can result in more coincidence between vectors attributed to samples and one in five class labels in an evaluation of 27 cancer types using RNA-seq data retrieved from TCGA. Enrichment in genes selected as expressed distinctly among class labels are also improved. Furthermore, it was found that the consideration of the strength of PPI as well as the restriction of genes to intersect between PPI and GE can drastically improve the coincidence.

## Author Contributions

Y.-H.T. planned the research and performed the analyses. Y.-H.T. and T.T. evaluated the results, discussions, and outcomes and wrote and reviewed the manuscript and has read and agreed to the published version of the manuscript.

## Data Availability Statement

All data analuysed in this paper can be available in GEO

## Disclaimer/Publisher’s Note

The statements, opinions and data contained in all publications are solely those of the individual author(s) and contributor(s) and not of MDPI and/or the editor(s). MDPI and/or the editor(s) disclaim responsibility for any injury to people or property resulting from any ideas, methods, instructions or products referred to in the content.

## References

1. Jalili, M.; Gebhardt, T.; Wolkenhauer, O.; Salehzadeh-Yazdi, A. Unveiling network-based functional features through integration of gene expression into protein networks. Biochimica et Biophysica Acta (BBA) - Molecular Basis of Disease 2018, 1864, 2349–2359. Accelerating Precision Medicine through Genetic and Genomic Big Data Analysis, https://doi.org/https://doi.org/10.1016/j.bbadis.2018.02.010.

2. Elbashir, M.K.; Mohammed, M.; Mwambi, H.; Omolo, B. Identification of Hub Genes Associated with Breast Cancer Using Integrated Gene Expression Data with Protein-Protein Interaction Network. Applied Sciences 2023, 13. https://doi.org/10.3390/app13042403.

3. Karimizadeh, E.; Sharifi-Zarchi, A.; Nikaein, H.; Salehi, S.; Salamatian, B.; Elmi, N.; Gharibdoost, F.; Mahmoudi, M. Analysis of gene expression profiles and protein-protein interaction networks in multiple tissues of systemic sclerosis. BMC Medical Genomics 2019, 12, 199. https://doi.org/10.1186/s12920-019-0632-2.

4. Tian, L.; Chen, T.; Lu, J.; Yan, J.; Zhang, Y.; Qin, P.; Ding, S.; Zhou, Y. Integrated Protein-Protein Interaction and Weighted Gene Co-expression Network Analysis Uncover Three Key Genes in Hepatoblastoma. Frontiers in Cell and Developmental Biology 2021, 9. https://doi.org/10.3389/fcell.2021.631982.

5. Wu, C.; Zhu, J.; Zhang, X. Integrating gene expression and protein-protein interaction network to prioritize cancer-associated genes. BMC Bioinformatics 2012, 13, 182. https://doi.org/10.1186/1471-2105-13-182.

6. Ewing, R.M.; Chu, P.; Elisma, F.; Li, H.; Taylor, P.; Climie, S.; McBroom-Cerajewski, L.; Robinson, M.D.; O’Connor, L.; Li, M.; et al. Large-scale mapping of human protein-protein interactions by mass spectrometry. Molecular Systems Biology 2007, 3, 89, [https://www.embopress.org/doi/pdf/10.1038/msb4100134]. https://doi.org/https://doi.org/10.1038/msb4100134.

7. Su, L.; Liu, G.; Guo, Y.; Zhang, X.; Zhu, X.; Wang, J. Integration of Protein-Protein Interaction Networks and Gene Expression Profiles Helps Detect Pancreatic Adenocarcinoma Candidate Genes. Frontiers in Genetics 2022, 13. https://doi.org/10.3389/fgene.2022.854661.

8. Zhong, J.; Tang, C.; Peng, W.; Xie, M.; Sun, Y.; Tang, Q.; Xiao, Q.; Yang, J. A novel essential protein identification method based on PPI networks and gene expression data. BMC Bioinformatics 2021, 22, 248. https://doi.org/10.1186/s12859-021-04175-8.

9. Taguchi, Y.H. Unsupervised Feature Extraction Applied to Bioinformatics; Springer International Publishing, 2020. https://doi.org/10.1007/978-3-030-22456-1.

10. Taguchi, Y.h.; Turki, T. Adapted tensor decomposition and PCA based unsupervised feature extraction select more biologically reasonable differentially expressed genes than conventional methods. Scientific Reports 2022, 12, 17438. https://doi.org/10.1038/s41598-022-21474-z.

11. Taguchi, Y.H.; Turki, T. TDbasedUFE and TDbasedUFEadv: bioconductor packages to perform tensor decomposition based unsupervised feature extraction. bioRxiv 2023, [https://www.biorxiv.org/content/early/2023/05/14/2023.05.14.540687.full.pdf]. https://doi.org/10.1101/2023.05.14.540687.

12. Nakerekanti, M.; Narasimha, V. Analysis on Malware Issues in Online Social Networking Sites (SNS). In Proceedings of the 2019 5th International Conference on Advanced Computing & Communication Systems (ICACCS), 2019, pp. 335–338. https://doi.org/10.1109/ICACCS.2019.8728536.

13. Xie, Z.; Bailey, A.; Kuleshov, M.V.; Clarke, D.J.B.; Evangelista, J.E.; Jenkins, S.L.; Lachmann, A.; Wojciechowicz, M.L.; Kropiwnicki, E.; Jagodnik, K.M.; et al. Gene Set Knowledge Discovery with Enrichr. Current Protocols 2021, 1, e90. [https://currentprotocols.onlinelibrary.wiley.com/doi/pdf/10.1002/cpz1.90]. https://doi.org/https://doi.org/10.1002/cpz1.90.

14. Jawaid, W. enrichR: Provides an R Interface to ‘Enrichr’, 2023. R package version 3.2.

15. Human protein-protein interaction network, 2018. https://snap.stanford.edu/biodata/datasets/10000/10000-PP-Pathways.html.

16. Oughtred, R.; Rust, J.; Chang, C.; Breitkreutz, B.J.; Stark, C.; Willems, A.; Boucher, L.; Leung, G.; Kolas, N.; Zhang, F.; et al. The BioGRID database: A comprehensive biomedical resource of curated protein, genetic, and chemical interactions. Protein Science 2021, 30, 187–200, [https://onlinelibrary.wiley.com/doi/pdf/10.1002/pro.3978]. https://doi.org/https://doi.org/10.1002/pro.3978.

17. Kosinski, M. RTCGA: The Cancer Genome Atlas Data Integration, 2022. R package version 1.28.0.

18. Kosinski, M. RTCGA.rnaseq: Rna-seq datasets from The Cancer Genome Atlas Project, 2022. R package version 20151101.28.0.

19. Kosinski, M. RTCGA.clinical: Clinical datasets from The Cancer Genome Atlas Project, 2022. R package version 20151101.28.0.

20. Brooks, A.J.; Putoczki, T. JAK-STAT Signalling Pathway in Cancer. Cancers 2020, 12. https://doi.org/10.3390/cancers12071971.

21. Lee, M.; Rhee, I. Cytokine Signaling in Tumor Progression. Immune Network 2017, 17, 214. https://doi.org/10.4110/in.2017.17.4.214.

22. Yang, J.; Nie, J.; Ma, X.; Wei, Y.; Peng, Y.; Wei, X. Targeting PI3K in cancer: mechanisms and advances in clinical trials. Molecular Cancer 2019, 18, 26. https://doi.org/10.1186/s12943-019-0954-x.

23. Yan, H.; Kamiya, T.; Suabjakyong, P.; Tsuji, N.M. Targeting C-Type Lectin Receptors for Cancer Immunity. Frontiers in Immunology 2015, 6. https://doi.org/10.3389/fimmu.2015.00408.

24. Mughini-Gras, L.; Schaapveld, M.; Kramers, J.; Mooij, S.; Neefjes-Borst, E.A.; Pelt, W.v.; Neefjes, J. Increased colon cancer risk after severe Salmonella infection. PLOS ONE 2018, 13, 1–19. https://doi.org/10.1371/journal.pone.0189721.

25. Wong, R.S. Apoptosis in cancer: from pathogenesis to treatment. Journal of Experimental & Clinical Cancer Research 2011, 30, 87. https://doi.org/10.1186/1756-9966-30-87.

26. Mantovani, A.; Petracca, G.; Beatrice, G.; Csermely, A.; Tilg, H.; Byrne, C.D.; Targher, G. Non-alcoholic fatty liver disease and increased risk of incident extrahepatic cancers: a meta-analysis of observational cohort studies. Gut 2022, 71, 778–788, [https://gut.bmj.com/content/71/4/778.full.pdf]. https://doi.org/10.1136/gutjnl-2021-324191.

27. Li, J.; Zhang, D.; Sun, Z.; Bai, C.; Zhao, L. Influenza in hospitalised patients with malignancy: a propensity score matching analysis. ESMO Open 2020, 5, e000968. https://doi.org/10.1136/esmoopen-2020-000968.

