## Supplementary material for "Integrated analysis of gene expression and protein-protein interaction with tensor decomposition": sample.R

```
require(Matrix)
# ----- uncomment the following three lines  only for Stanford PPI
x <- read.csv("PP-Pathways_ppi.csv.gz",sep=",",header=F)
gene <- unique(c(x[,1],x[,2]))
xx <- cbind(match(x[,1],gene),match(x[,2],gene))

# ----- uncomment the following three lines  only  for BIOGRID PPI
#x <- read.csv(unz("BIOGRID-MV-Physical-4.4.221.tab2.zip", "BIOGRID-MV-Physical-4.4.221.tab2.txt"), header = TRUE,sep = "\t")
#gene <- unique(c(x[,2],x[,3]))
#xx <- cbind(match(x[,2],gene),match(x[,3],gene))

xx <- xx[xx[,1]!=xx[,2],]
X <- sparseMatrix(c(xx[,1],xx[,2]),c(xx[,2],xx[,1]),x=1)
X <- X+t(X)
require(RTCGA)
require(RTCGA.rnaseq)
require(RTCGA.clinical)
gene1 <- unlist(lapply(strsplit(colnames(ACC.rnaseq),"|",fixed=T),"[",2))
gene_int <- intersect(gene1,gene)
index0 <- match(gene,gene_int)
index0 <- index0[!is.na(index0)]
XX <- X[index0,index0]
index <- match(gene1,gene_int)
index <- index[!is.na(index)]

require(irlba)
require(rTensor)
SVD <- irlba(XX,10)

require(RTCGA.clinical)
LIST1 <- list(ACC.clinical,
              BLCA.clinical,
              BRCA.clinical,
              CESC.clinical,
              COAD.clinical,
              ESCA.clinical,
              GBM.clinical,
              HNSC.clinical,
              KICH.clinical,
              KIRC.clinical,
              KIRP.clinical,
              LGG.clinical,
              LIHC.clinical,
              LUAD.clinical,
              LUSC.clinical,
              OV.clinical,
              PAAD.clinical,
              PCPG.clinical,
              PRAD.clinical,
              READ.clinical,
              SARC.clinical,
              SKCM.clinical,
              STAD.clinical,
              TGCT.clinical,
              THCA.clinical,
              UCEC.clinical,
              UCS.clinical)

require(RTCGA.rnaseq)
LIST <- list(ACC.rnaseq,
             BLCA.rnaseq,
             BRCA.rnaseq,
             CESC.rnaseq,
             COAD.rnaseq,
             ESCA.rnaseq,
             GBM.rnaseq,
             HNSC.rnaseq,
             KICH.rnaseq,
             KIRC.rnaseq,
             KIRP.rnaseq,
             LGG.rnaseq,
             LIHC.rnaseq,
             LUAD.rnaseq,
             LUSC.rnaseq,
             OV.rnaseq,
             PAAD.rnaseq,
             PCPG.rnaseq,
             PRAD.rnaseq,
             READ.rnaseq,
             SARC.rnaseq,
             SKCM.rnaseq,
             STAD.rnaseq,
             TGCT.rnaseq,
             THCA.rnaseq,
             UCEC.rnaseq,
             UCS.rnaseq)

P_all <- array(NA,c(10,3,length(LIST)))
for (i in seq_len(length(LIST)))
{
  cat(i," ")
  gid <- grep("patient.bcr_patient_barcode",colnames(LIST1[[i]]))   
  index1 <- match(tolower(substring(LIST[[i]][,1],1,12)),
                  LIST1[[i]][,gid])
  XX1 <- data.matrix(LIST[[i]][,index])
  SVD1 <- svd(XX1,10)
  #----- uncomment one of the following two lines
  Z <- array(NA,c(16774,10,2)) #stanford
  #Z <- array(NA,c(11294,10,2)) #BIOGRID
  #------------------------------------
  Z[,,1] <- SVD$v[,1:10]
  Z[,,2] <- SVD1$v[,1:10]
  HOSVD1 <- hosvd(as.tensor(Z),c(10,10,2))
  #------ uncomment one of class labels ----
  #k<- grep("patient.stage_event.pathologic_stage",colnames(LIST1[[i]]))
  k<- grep("patient.stage_event.tnm_categories.pathologic_categories.pathologic_m",colnames(LIST1[[i]]))
  #k<- grep("patient.stage_event.tnm_categories.pathologic_categories.pathologic_n",colnames(LIST1[[i]]))
  #k<- grep("patient.stage_event.tnm_categories.pathologic_categories.pathologic_t",colnames(LIST1[[i]]))
  #k<- grep("patient.vital_status",colnames(LIST1[[i]]))
  #---------------------------------------
  label <- LIST1[[i]][index1,k]
  if (sum(!is.na(label))!=0)
  {
    P<-NULL
    for (j in seq_len(3))
    {
      if (j==1) {LM <- lm(XX1 %*% SVD1$v[,1:10]~label)} else if (j==2)
      {
        LM <- lm(XX1 %*% HOSVD1$U[[1]]~label)
      } else {
        LM <- lm(XX1 %*% SVD$v[,1:10]~label)
      } 
      SLM <- summary(LM)
      fs <- t(data.frame(lapply(SLM,"[",10)))
      P <- cbind(P,pf(fs[,1],fs[,2],fs[,3],lower.tail=F))
    }
    P_all[,,i] <-P
  } else {
    P_all[,,i] <-NA
  }
  
}
ina <- apply(is.na(P_all),3,sum)
#P_all[l,j,i] 
#l: lth singular value vectors. l is less than or equal to 10
#j: 1 -> SVD, 2 -> HOSVD, 3 -> simple integration
#i: one of 27 cancer types. i is less than or equal to 27
#Only the following is used fot t test and Wilcox test, since ina==0 means no missng values
P_all[, j, ina == 0]


#enrichment analysis
options(timeout = 4000000) 
require(mygene)
library(enrichR)
enriched_all <- rep(list(NA),length(LIST))
for (i in seq_len(length(LIST)))
{
  cat(i," \n")
  gid <- grep("patient.bcr_patient_barcode",colnames(LIST1[[i]]))   
  index1 <- match(tolower(substring(LIST[[i]][,1],1,12)),
                  LIST1[[i]][,gid])
  XX1 <- data.matrix(LIST[[i]][,index])
  SVD1 <- svd(XX1,10)
  #----- uncomment one of the following two lines
  Z <- array(NA,c(16774,10,2)) #stanford
  #Z <- array(NA,c(11294,10,2)) #BIOGRID
  #-----------------------------
  Z[,,1] <- SVD$v[,1:10]
  Z[,,2] <- SVD1$v[,1:10]
  #uncomment only for HOSVD
  #HOSVD1 <- hosvd(as.tensor(Z),c(10,10,2)) #HOSVD
  #k<- grep("patient.stage_event.pathologic_stage",colnames(LIST1[[i]]))
  #k<- grep("patient.stage_event.tnm_categories.pathologic_categories.pathologic_m",colnames(LIST1[[i]]))
  #k<- grep("patient.stage_event.tnm_categories.pathologic_categories.pathologic_n",colnames(LIST1[[i]]))
  k<- grep("patient.stage_event.tnm_categories.pathologic_categories.pathologic_t",colnames(LIST1[[i]]))
  #k<- grep("patient.vital_status",colnames(LIST1[[i]]))
  label <- LIST1[[i]][index1,k]
  if (sum(!is.na(label))!=0)
  {
    #----- uncomment one of the following two lines
    #LM <- lm(XX1 %*% HOSVD1$U[[1]]~label) #HOSVD
    LM <- lm(XX1 %*% SVD1$v[,1:10]~label) #SVD
    #----------------------------------
    SLM <- summary(LM)
    fs <- t(data.frame(lapply(SLM,"[",10)))
    P <- pf(fs[,1],fs[,2],fs[,3],lower.tail=F)
      th <- function(sd){
      P2 <- pchisq((HOSVD1$U[[1]][,l]/sd)^2,1,lower.tail=F)
      hc<- hist(1-P2,breaks=100,plot=F)
      return(sd(hc$count[1:sum(hc$breaks<1-min(P2[p.adjust(P2,"BH")>0.01]))]))
    }
    l <- which.min(P)
    sd <- optim(1e-4,th)$par
    P2 <- pchisq((HOSVD1$U[[1]][,l]/sd)^2,1,lower.tail=F)
    gene_sel <- gene_int[index0][p.adjust(P2,"BH")<0.01]
    result <- getGenes(geneid = gene_sel, fields = c("symbol"))
    gene_symbol<- result$symbol
    setEnrichrSite("Enrichr")
    websiteLive <- TRUE
    dbs <- c("KEGG_2021_Human",
             "GO_Biological_Process_2023", "GO_Cellular_Component_2023",
             "GO_Molecular_Function_2023"
    )
    enriched_all[[i]] <- enrichr(gene_symbol, dbs)
  } else {
    enriched_all[[i]] <- NA
  }
} 
```
