## Supplementary material for "Integrated analysis of gene expression and protein-protein interaction with tensor decomposition": Supplementary_Tables.pdf

Table S1: Numerical values of bar plots shown in Fig. 2

| | "vital_status" | "pathologic_stage" | "pathologic_m" | "pathologic_t" | "pathologic_n" |
|---|---|---|---|---|---|
| | Stanford | | | | |
| Wilcoxon test | $2.14 \times 10^{-8}$ | $3.18 \times 10^{-18}$ | $6.69 \times 10^{-1}$ | $6.43 \times 10^{-22}$ | $8.67 \times 10^{-1}$ |
| | BIOGRID | | | | |
| Wilcoxon test | $1.03 \times 10^{-30}$ | $1.34 \times 10^{-5}$ | $3.16 \times 10^{-15}$ | $8.53 \times 10^{-29}$ | $3.93 \times 10^{-18}$ |

Table S2: Numerical values of bar plots shown in Fig. 10

| | "vital_status" | "pathologic_stage" | "pathologic_m" | "pathologic_t" | "pathologic_n" |
|---|---|---|---|---|---|
| | Correlation coefficients | | | | |
| Stanford | 0.531493 | 0.5631397 | 0.4323562 | 0.3987178 | 0.5888728 |
| BIOGRID | 0.5604814 | 0.5807142 | 0.6080881 | 0.6166185 | 0.5630905 |
| | $P$-values | | | | |
| Stanford | $4.36 \times 10^{-21}$ | $1.89 \times 10^{-16}$ | $1.35 \times 10^{-9}$ | $4.98 \times 10^{-9}$ | $4.68 \times 10^{-20}$ |
| BIOGRID | $9.80 \times 10^{-24}$ | $1.29 \times 10^{-17}$ | $1.39 \times 10^{-19}$ | $2.46 \times 10^{-22}$ | $3.99 \times 10^{-18}$ |

Table S3: Numerical values of bar plots shown in Fig. 11

| | "vital_status" | "pathologic_stage" | "pathologic_m" | "pathologic_t" | "pathologic_n" |
|---|---|---|---|---|---|
| | Stanford | | | | |
| Wilcoxon test | 1 | 1 | 0.991 | $4.22 \times 10^{-19}$ | 1 |
| | BIOGRID | | | | |
| Wilcoxon test | 1 | 1 | $9.23 \times 10^{-17}$ | 1 | 1 |
