## Supplementary figures and images for "Integrated analysis of gene expression and protein-protein interaction with tensor decomposition"

### enrihed_all.pdf

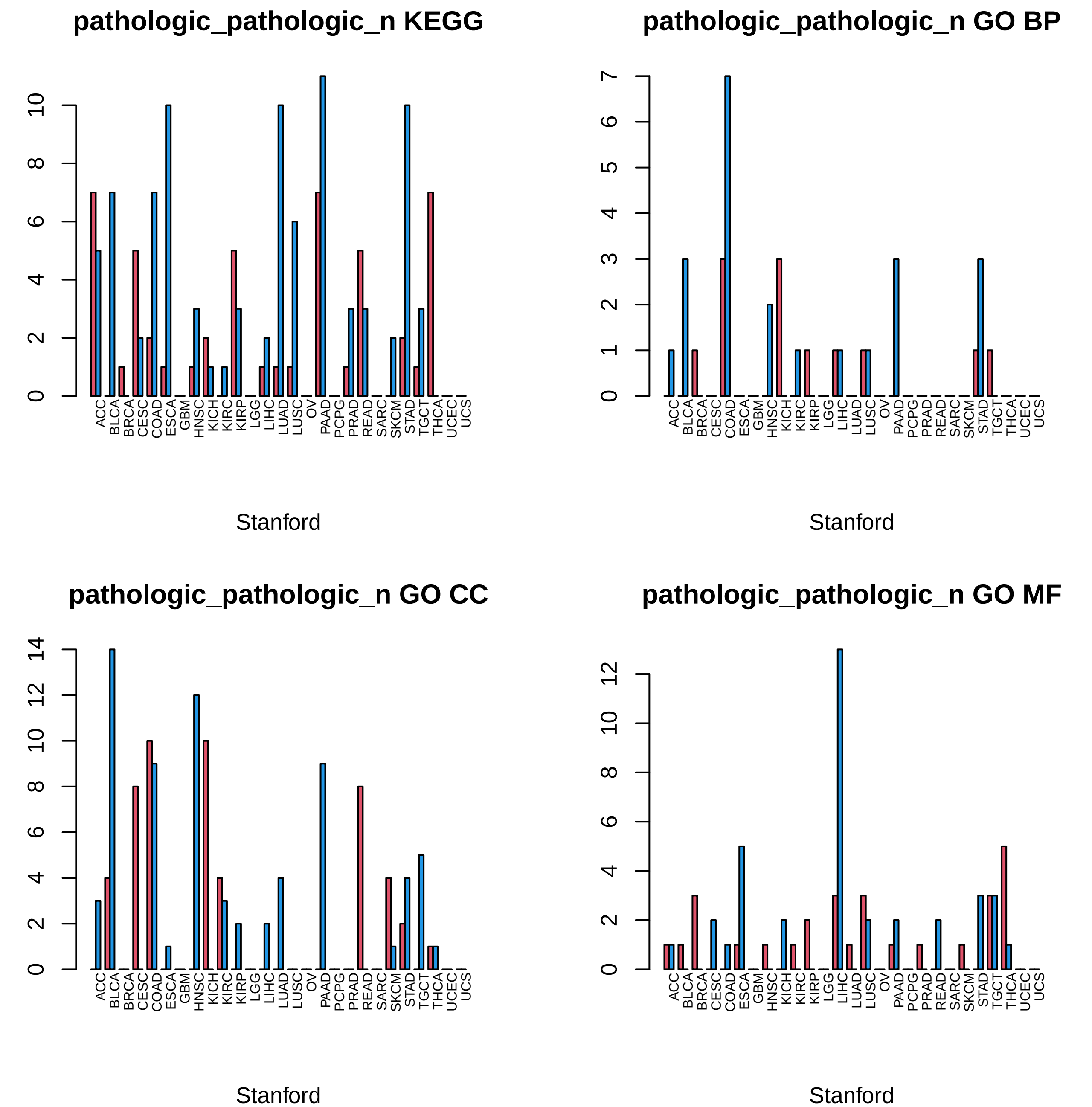

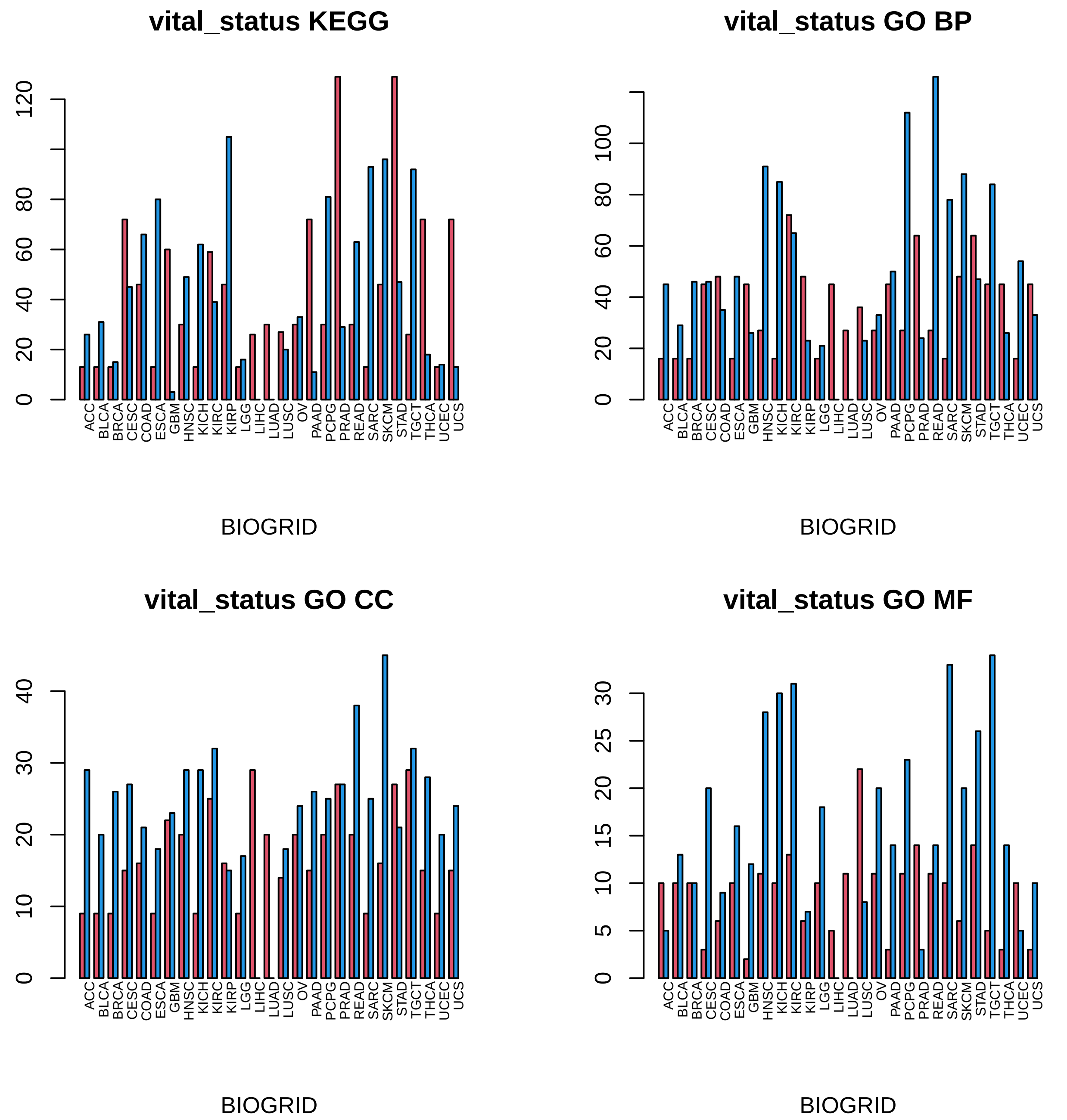

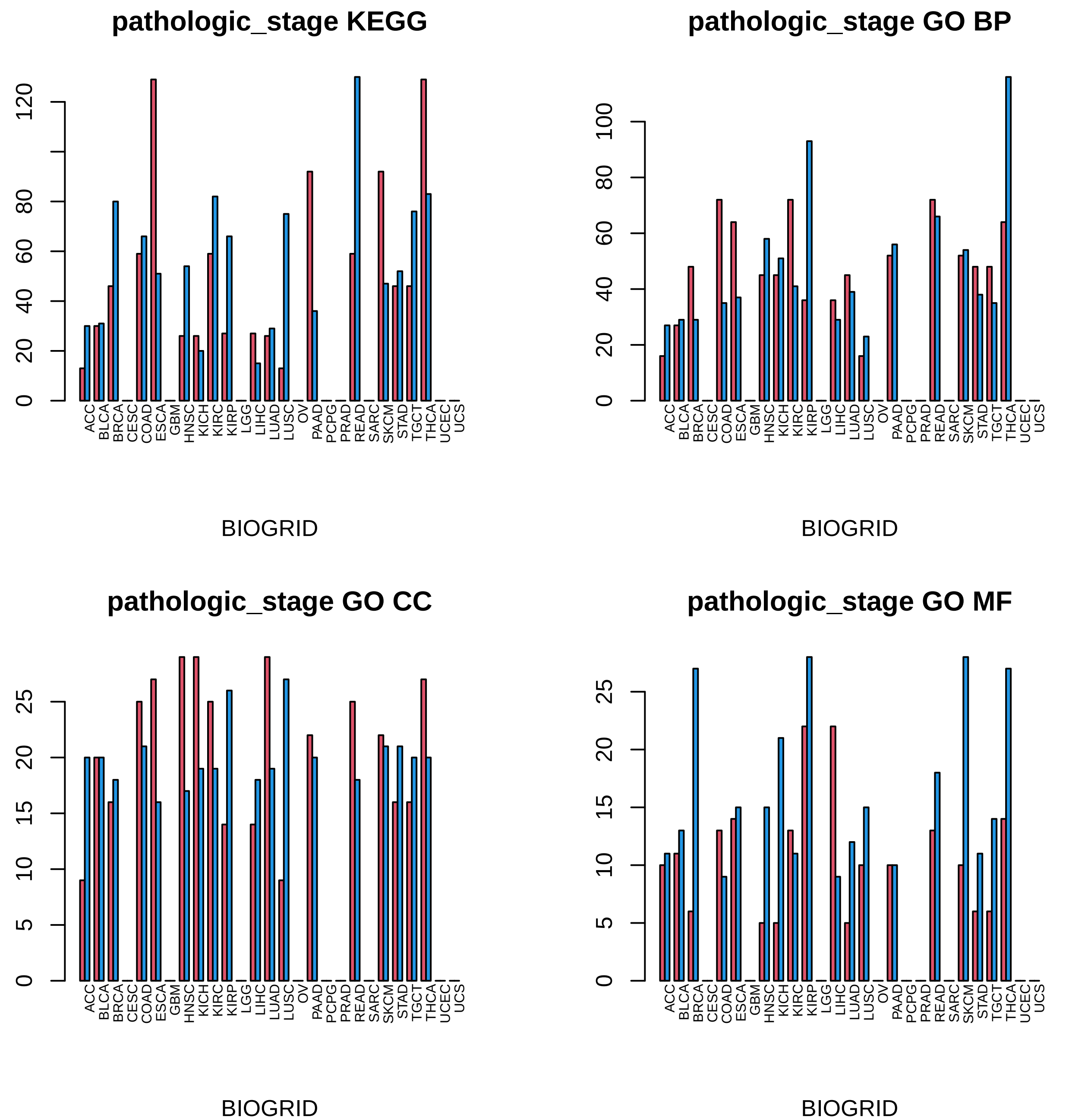

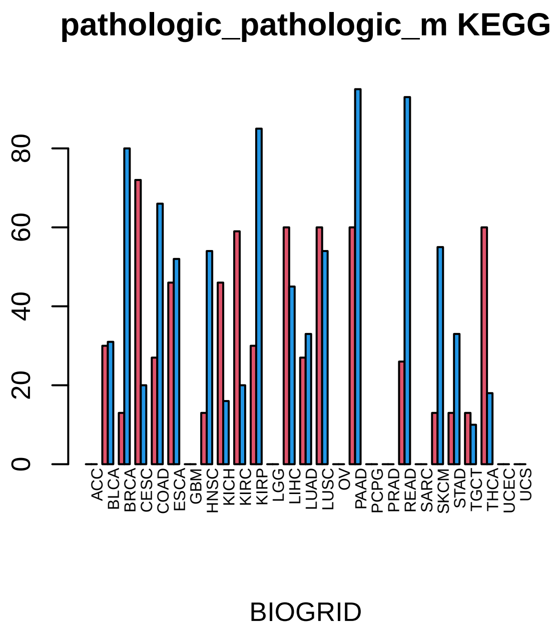

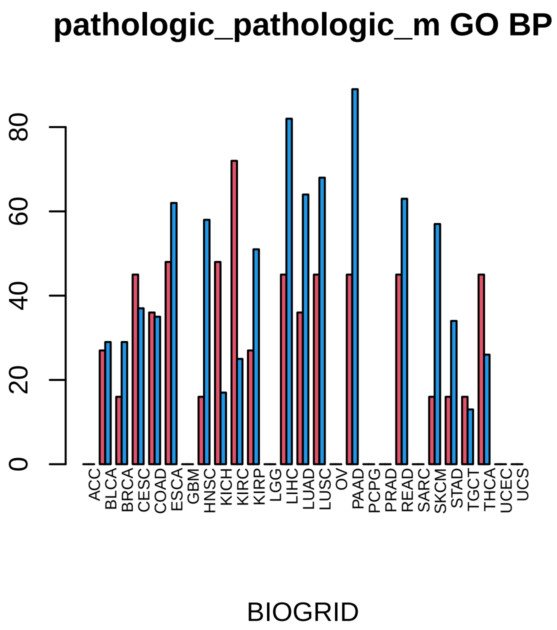

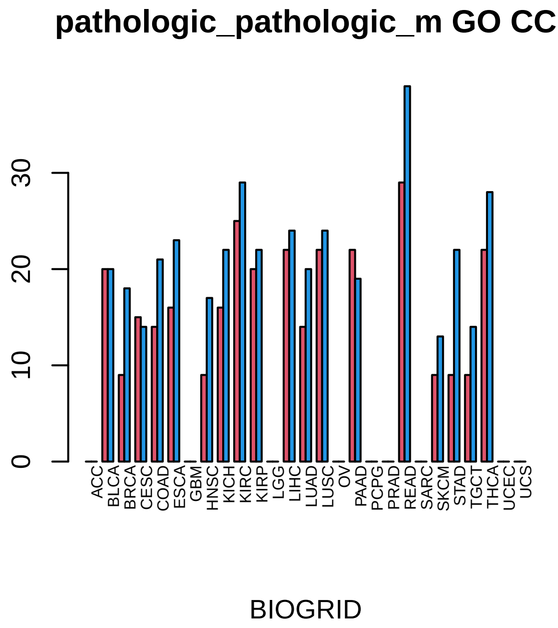

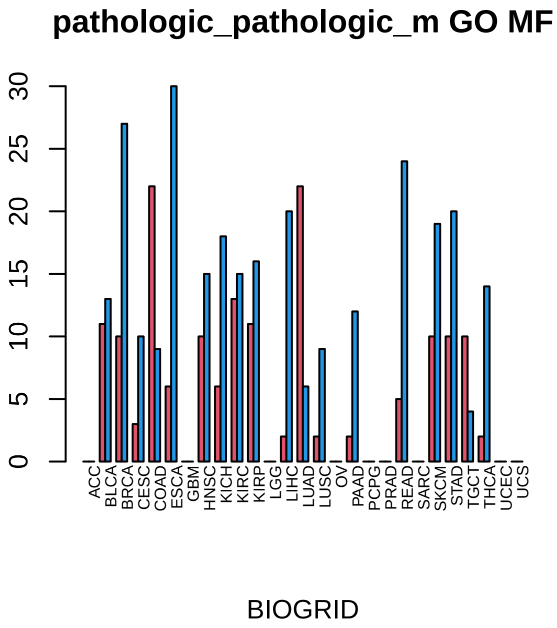

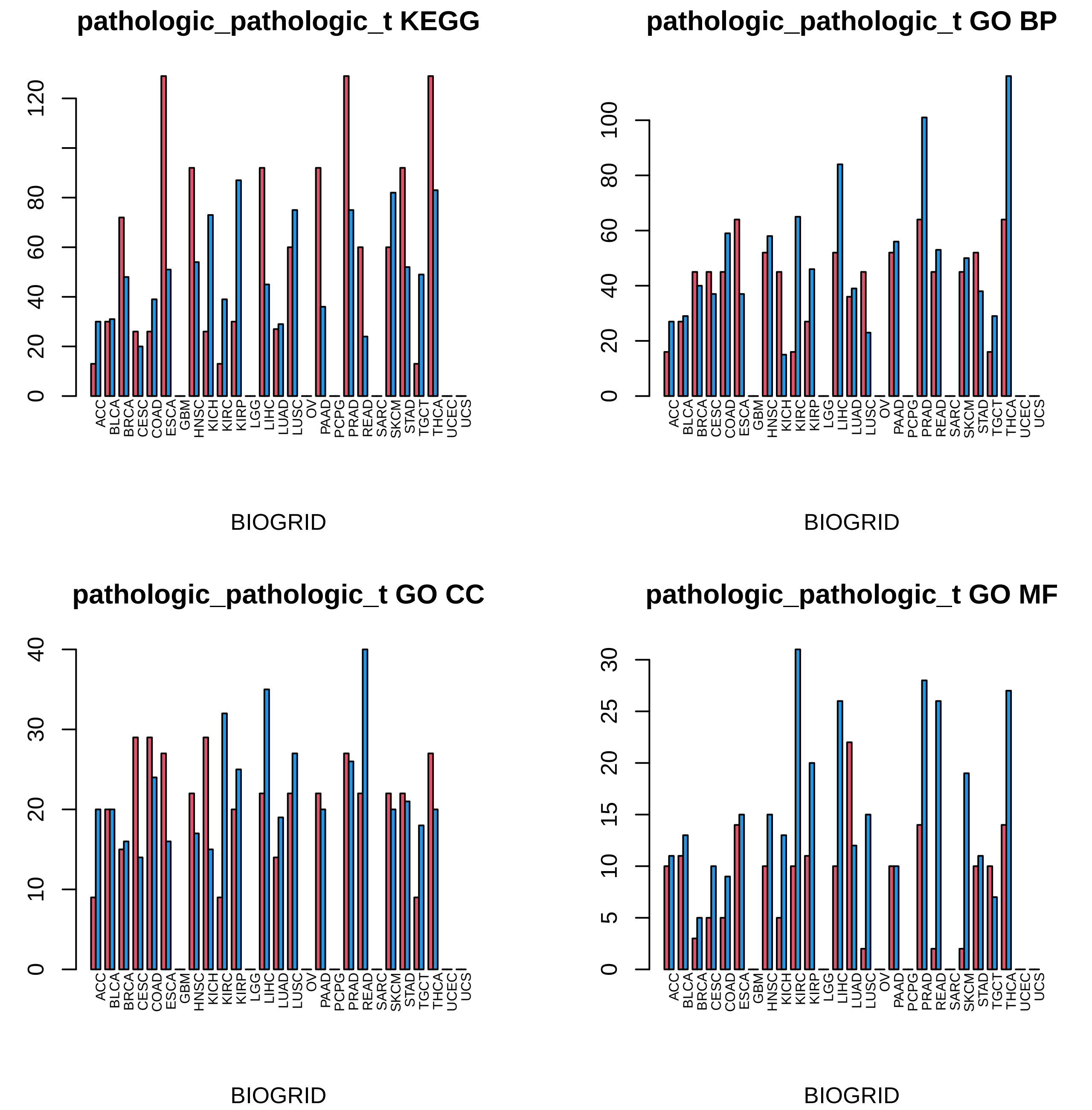

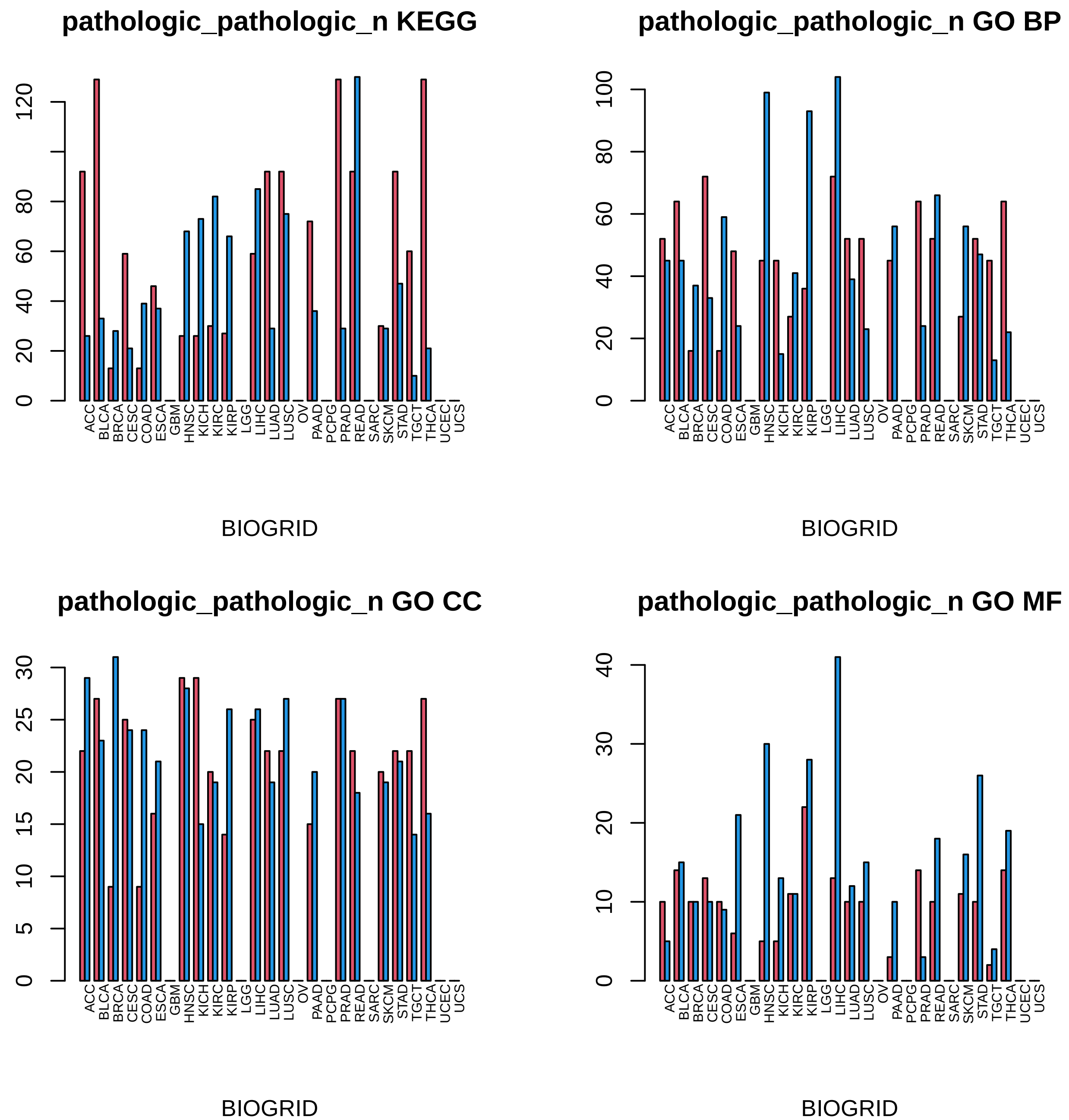
